# Sulfur oxidation and reduction are coupled to nitrogen fixation in the roots of a salt marsh foundation plant species

**DOI:** 10.1101/2023.05.01.538948

**Authors:** J.L. Rolando, M. Kolton, T. Song, Y. Liu, P. Pinamang, R. Conrad, J.T. Morris, K.T. Konstantinidis, J.E. Kostka

## Abstract

Symbiotic root microbiota are crucial for plant growth as they assist their hosts in nutrient acquisition. In the roots of coastal marine plants, heterotrophic activity in the rhizosphere by sulfate-reducing microorganisms has been linked to nitrogen fixation. In this study, we recovered 239 high-quality metagenome-assembled genomes (MAGs) from a salt marsh dominated by the foundation plant *Spartina alterniflora*, including diazotrophic sulfate-reducing and sulfur-oxidizing bacteria thriving in the root compartment. Here we show for the first time that highly-abundant sulfur-oxidizing bacteria in the roots of a coastal macrophyte encode and highly express genes for nitrogen fixation (nifHDK). Further, we leveraged a *S. alterniflora* biomass gradient to gain a mechanistic understanding on how root-microbe interactions respond to abiotic stress from anoxia and elevated sulfide concentration. We observed that the roots of the stressed *S. alterniflora* phenotype exhibited the highest rates of nitrogen fixation and expression levels of both the oxidative and reductive forms of the dissimilatory sulfite reductase gene (dsrAB). Approximately 25% and 15% of all sulfur-oxidizing dsrA and nitrogen-fixing nifK transcripts, respectively, were associated with novel MAGs of the *Candidatus* Thiodiazotropha genus in the roots of the stressed *S. alterniflora* phenotype. We conclude that the rapid cycling of sulfur in the dynamic *S. alterniflora* root zone is coupled to nitrogen fixation during both reductive and oxidative sulfur reactions, and that the *S. alterniflora* – *Ca.* Thiodiazotropha symbiosis is an adaptive response to anoxic and sulfidic sediment conditions, whereby the plants benefit from reduced sulfide toxicity and potential nitrogen acquisition.

## Introduction

The microbial communities closely associated with plant hosts (i.e., plant microbiota) have received substantial attention due to their potential role in plant nutrient acquisition, phytohormone synthesis, prevention of soil-borne disease, and detoxification of the rhizosphere and root environment (Bulgarelli et al., 2013; Trivedi et al., 2020). Most plant microbiome studies have been performed in terrestrial ecosystems with an emphasis on agricultural plants (Trivedi et al., 2020). Plant microbiota from vegetated coastal ecosystems (i.e., seagrass meadows, mangroves, and salt marshes) remain mostly understudied, even though they play a key role in global climate regulation and the cycling of major nutrients (Barbier et al., 2011; Duarte et al., 2013). Covering less than 3% of the global surface area occupied by forests, coastal vegetated ecosystems exhibit an equivalent rate of carbon sequestration at the planetary scale due to their excess in primary production over organic matter decomposition (Duarte et al., 2013). Further, vegetated estuarine ecosystems play a key role in water purification by reducing coastal nutrient pollution (Barbier et al., 2011). Despite their high ecological value, little is known about how plant-microbe interactions contribute to the functioning of coastal marine ecosystems, their resilience to climate change, and provisioning of ecosystem services.

Previous studies have shown that the root zone of coastal marine plants is a hotspot for the cycling of carbon, nitrogen, and sulfur (Gandy and Yoch, 1988; Welsh et al., 1996; Spivak and Reeve, 2015; Thomas et al., 2014; Crump et al., 2018; Kolton et al., 2020). In coastal Georgia, US, sulfate-reducing and sulfur-oxidizing bacteria were recently shown to be overrepresented in the core root microbiome of the salt marsh foundation plant *Spartina alterniflora*, comprised of persistent taxa across multiple plant individuals (Rolando et al., 2022). Sulfate reduction, an anaerobic respiration pathway, represents a dominant terminal electron-accepting process coupled to the breakdown of organic matter in marine ecosystems (Kostka et al., 2002; Jørgensen et al., 2019). Evidence from experimental and field research indicates that rhizospheric sulfate-reducing bacteria assimilate photosynthates from *S. alterniflora*, while fueling nitrogen fixation (Gandy and Yoch, 1988; Spivak and Reeve, 2015). Similarly, rates of nitrogen fixation have been closely associated with organic matter degradation by sulfate reduction in seagrass meadows (McGlathery et al., 1998; Herbert, 1999; Nielsen et al., 2001; Welsh et al., 1996). Conversely, the oxidation of reduced sulfur compounds is a chemolithotrophic process requiring potent terminal electron acceptors such as oxygen, nitrate, or oxidized metals. In particular, sulfide is a known phytotoxin, and its belowground oxidation is considered a detoxifying reaction for plants inhabiting coastal marine ecosystems (Lamers et al., 2013). Based on amplicon sequencing of the 16S rRNA gene, sulfur oxidation coupled with nitrogen fixation was recently hypothesized as an important process for plant growth under sulfidic conditions (Rolando et al., 2022). Nitrogen is often the limiting nutrient for salt marsh plants, and stress from sulfide toxicity and anoxia further impairs root active acquisition of nitrogen (Mendelssohn and Morris, 2000). Close relatives of diazotrophic sulfur-oxidizing symbionts from the *Sedimenticolaceae* family (genus: *Candidatus* Thiodiazotropha) have been shown to inhabit the roots of seagrasses and marsh plants in coastal marine ecosystems (Martin et al., 2020a, Rolando et al., 2022). The *Ca.* Thiodiazotropha genus was first discovered as bacterial symbionts of lucinid clams, where they provide both fixed carbon and nitrogen to their animal host by sulfur-mediated chemolithoautotrophy (Petersen et al., 2017). Most studies assessing the ecology and function of sulfur chemosymbiosis have been performed within marine invertebrates hosts (Petersen et al., 2017; Osvatic et al., 2021, 2023; Lim et al., 2019a, 2019b). In contrast, there is still no evidence for the coupling of sulfur oxidation with nitrogen and/or carbon fixation in the roots of coastal marine plants because the genomes, metabolic potential, and activity of root-associated sulfur-oxidizing bacteria are still lacking. Limited studies have looked into the composition and abundance of sulfur-oxidizing bacteria present in the roots of coastal marine plants. These studies mainly relied on microscopy- and DNA/RNA amplicon-based community analyses (Thomas et al., 2014; Martin et al., 2020a; Kolton et al., 2020; Rolando et al., 2022). Only one study utilized a metatranscriptomics approach to investigate the gene expression of seagrass root-associated microbiomes (Crump et al., 2018). However, genome-wide gene expression profiles of sulfur-oxidizing bacteria and the coupling of sulfur oxidation with other biogeochemical processes are still to be deciphered in the roots of coastal marine plants.

Ecosystem models and empirical evidence indicate that climate change is altering the hydrology, biogeochemistry, and plant community composition of coastal wetlands (Guimond et al., et al., 2020; Donnelly and Bertness, 2001). Thus, a mechanistic understanding of how environmental perturbations impact plant-microbe interactions will be critical to forecasting the resilience of salt marsh ecosystems to climate change. According to the stress-gradient hypothesis (SGH), the frequencies of facilitative and competitive interactions are inversely related along abiotic stress gradients, with facilitative interactions occurring more frequently under high abiotic stress than under benign conditions (Bertness and Callaway, 1994). This concept has mostly been explored in plant-plant interactions; whereas, limited studies assessing plant-microbe interactions have found that the SGH is consistent in certain plant phenological stages (David et al., 2020). Following the SGH, we hypothesize that beneficial plant-microbe interactions will be more common in stressed coastal marine ecosystems, where plants experience greater salinity, redox, and sulfide toxicity stress. *S. alterniflora*-dominated salt marsh ecosystems represent an ideal natural laboratory in which to study the effects of stress on plant-microbe interactions because steep gradients in plant productivity are formed within short distances (Mendelssohn and Morris, 2000). The productivity gradient is the result of decreased soil redox potential, along with increased salinity and anoxia, extending from vegetated tidal creek banks towards the interior of the marsh (Figure 1). Higher primary productivity is reflected in tall *S. alterniflora* plants (> 80 cm) growing adjacent to tidal creek banks, while smaller plants (< 50 cm) inhabit the interior of the marsh. We hypothesize that the strength of plant-sulfur chemosymbiosis increases in stressed marshes where elevated sulfide oxidation rates are coupled to nitrogen fixation.

**Figure 1.**
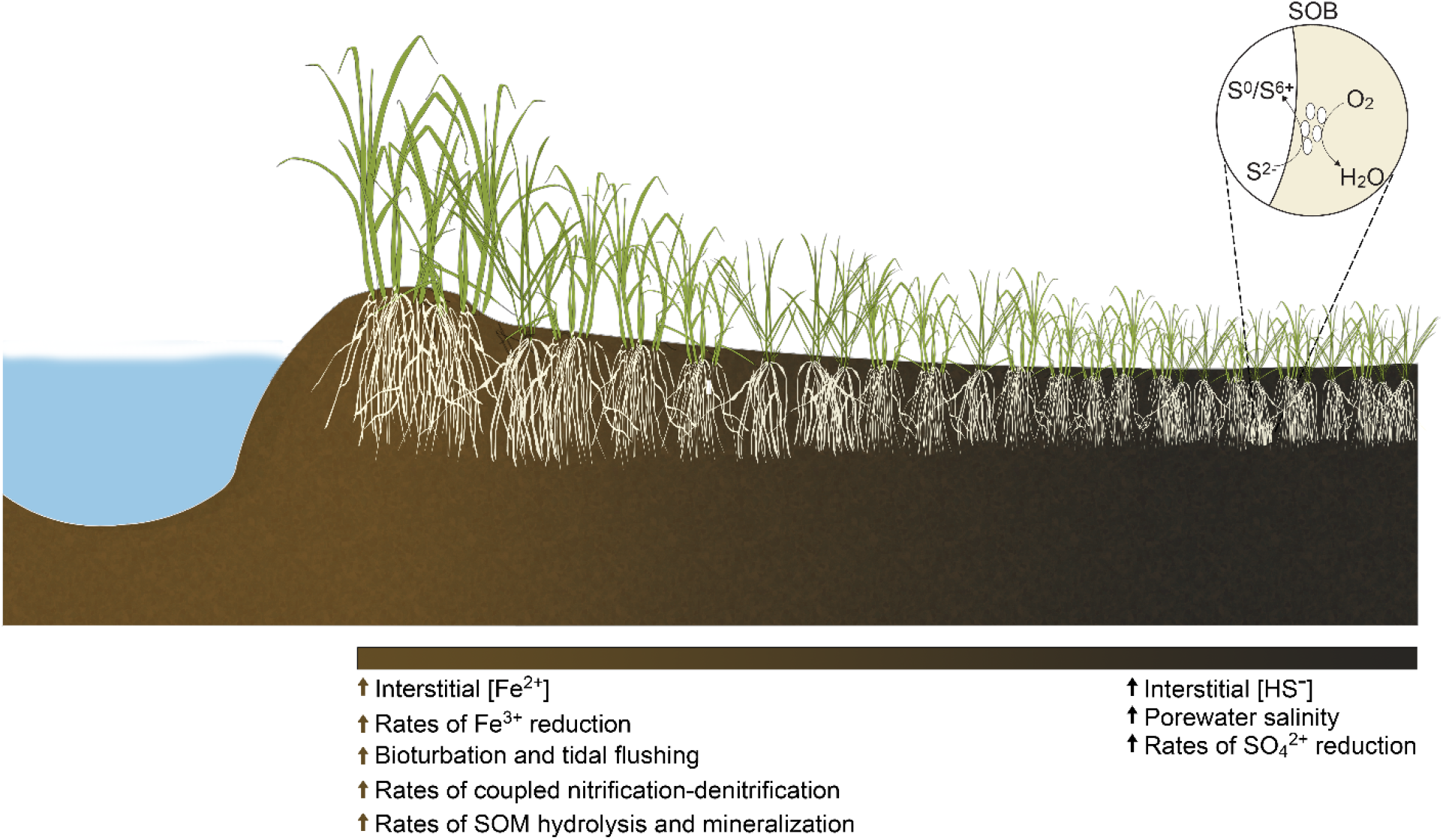
*Spartina alterniflora* biomass gradient as a natural laboratory. A gradient in *S. alterniflora* aboveground biomass is commonly observed with tall plants growing at the levees next to large tidal creeks and a short phenotype of *S. alterniflora* dominating the interior of the marsh. Sediments from the tall *S. alterniflora* zone are characterized as a more oxidized environment with higher levels of iron, coupled nitrification-denitrification as well as higher rates of organic matter hydrolysis and mineralization. Conversely, sediments from the short phenotype tend to be more chemically reduced, with higher rates of sulfate reduction, elevated porewater salinity, and less bioturbation and tidal flushing. Roots from the short *S. alterniflora* phenotype have been proposed to harbor sulfur-oxidizing bacteria (SOB) that benefit the plant by detoxifying the root environment.

Thus, the objectives of the present study were: (i.), to demonstrate the coupling of sulfur oxidation with nitrogen fixation in the root environment of *S. alterniflora* and (ii.) to evaluate the effect of environmental stress on the assembly, activity, and ecological interactions of the *S. alterniflora* root microbiome.

## 2. Results

### 2.1 Genome-centric multi-omics approach reveals prokaryotic sulfur metabolism is coupled to nitrogen fixation in the roots of a foundation salt marsh plant

We studied the root microbiome of the salt marsh foundation species of the US Atlantic and Gulf of Mexico coastlines, *Spartina alterniflora*, at Sapelo Island, GA during the summers of 2018, 2019 and 2020. A combination of prokaryotic DNA and RNA quantification, shotgun metagenomics, metatranscriptomics, and rate measurements of two key pathways of the nitrogen cycle were performed along a stress gradient within three compartments: sediment, rhizosphere and root (Figure 1). The short and tall *S. alterniflora* phenotypes represent the end extremes of the plant biomass gradient formed by stress from salinity, sulfide toxicity and anoxia (Figure 1). Four biological replicates per compartment and *S. alterniflora* phenotype were sequenced for both metagenomic and transcriptomic analysis (Further details about sequencing in Materials and Methods and Supplementary Table S1).

Using a custom-designed bioinformatics pipeline, we binned 239 high-quality metagenome assembled genomes (MAGs; > 50 Quality score). MAGs were dereplicated into 160 genomospecies (gsp, plural: gspp) by grouping them according to 95% average nucleotide identity (ANI) (Figure 2). MAGs were assigned to 19 phyla. Almost half of the recovered genomes were taxonomically affiliated with the *Proteobacteria* or *Desulfobacterota* phyla (Figure 2, further MAGs taxonomic and statistical information in Supplementary Table S2). Taxonomic novelty was assessed using GTDB-Tk v2.1.0 with the reference database R07-RS207 (Chaumeil et al., 2022). Recovered MAGs included 1 gspp from a previously undescribed order, 10 gspp from undescribed families, 52 gspp from undescribed genera, 94 gspp from undescribed species, and only 3 gspp from previously described species. In order to assess the MAGs’ genetic potential to perform important biogeochemical functions in the salt marsh environment, we annotated their open reading frames (ORFs) using the eggNOG database, and focused on predicted genes involved in the biogeochemical cycling of carbon, nitrogen, and sulfur (Supplementary Table S3).

**Figure 2.**
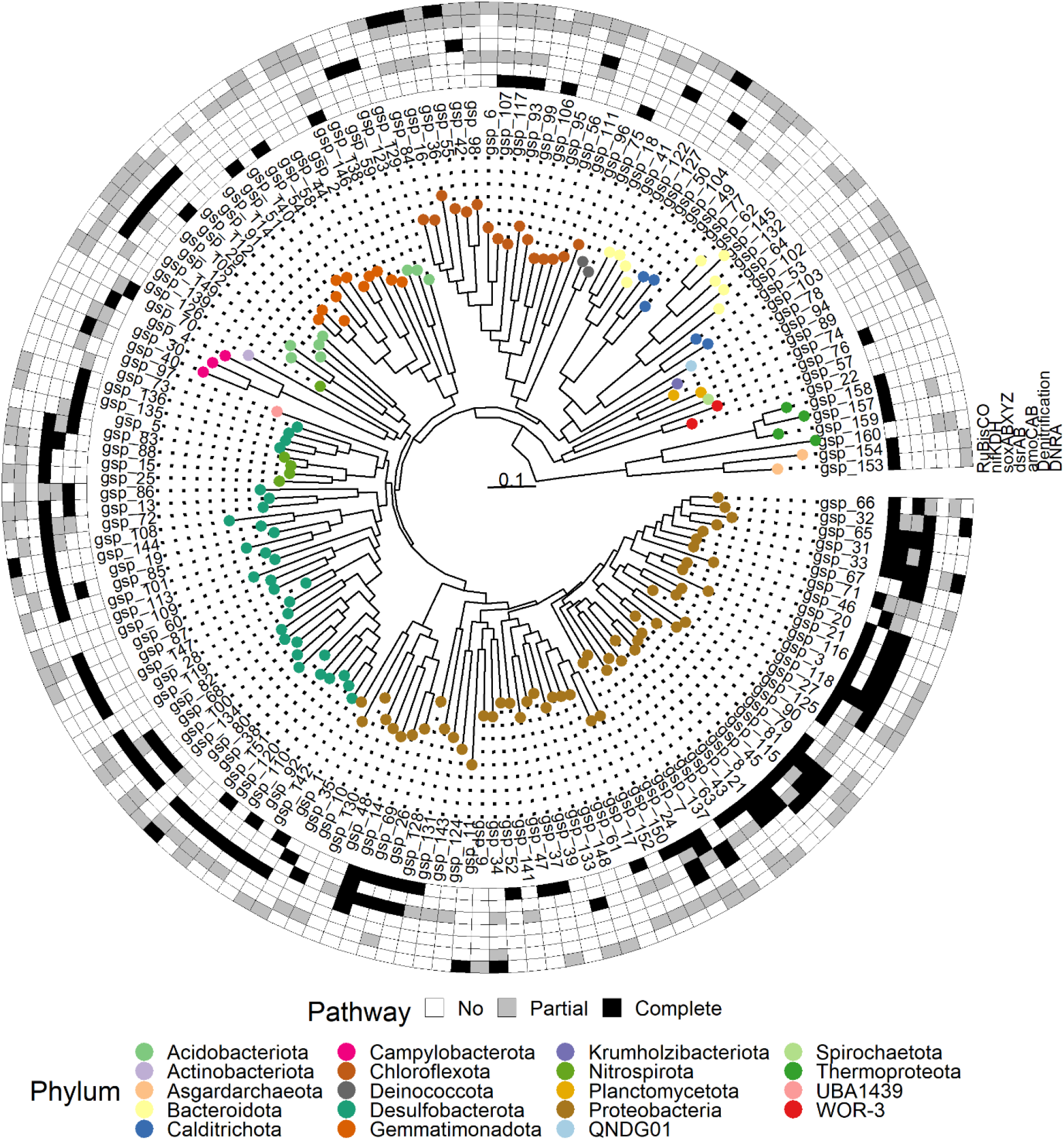
Phylogenetic reconstruction of 160 dereplicated high-quality metagenome assembled genomes (MAGs, > 50 quality-score) binned from *Spartina alterniflora* sediment, rhizosphere, and root samples. Outer rim shows the presence/absence of genes for carbon fixation (RuBisCO), nitrogen fixation (nifKDH), thiosulfate oxidation (soxABXYZ), dissimilatory sulfite reduction/oxidation (dsrAB), nitrification (amoCAB), denitrification, and dissimilatory nitrate reduction to ammonium (DNRA).

A large proportion of binned *Proteobacteria*, including members of the *Ca.* Thiodiazotropha genus, contained genes for nitrogen fixation, carbon fixation through RuBisCO, as well as for dissimilatory sulfur metabolism, and thiosulfate oxidation using the soxABXYZ complex (Figure 2). Most MAGs from the *Desulfobacterota* phylum presented genes for dissimilatory metabolism of sulfur, including members of the *Desulfosarcinaceae* family (Figure 2). Because the dissimilatory sulfite reductase enzyme (dsrAB) can function in either sulfur reduction or oxidation, we performed a phylogenetic analysis to identify the dsrAB type from our MAGs. We aligned recovered dsrAB genes to a reference alignment (Müller et al., 2015) and annotated the dsrAB type based on placement in an approximately-maximum-likelihood phylogenetic tree (Supplementary Figure S1). All dsrAB genes retrieved from *Proteobacteria* gspp (*Alpha*- and *Gammaproteobacteria* gspp) were placed within the oxidative type; while dsrAB genes recovered from *Acidobacteriota*, *Bacteroidota*, *Chloroflexota*, *Desulfobacterota*, *Gemmatimonadota*, and *Nitrospirota* gspp clustered within the reductive bacterial dsrAB type (Supplementary Figures S1, S2).

We measure rates of ^15^N_2_ fixation under oxic and anoxic conditions by sediment, a rhizosphere-root mix, and root samples. Rates were measured under oxic and anoxic conditions to capture the oxygen fluctuation experienced by microorganisms in the root-zone of *S. alterniflora*. The two *S. alterniflora* phenotypes exhibited significantly higher N fixation rates in their root tissue compared to the sediment or rhizosphere-root mix under both aerobic and anaerobic conditions (Figure 3). Root tissue from the stressed short *S. alterniflora* phenotype showed 1.9- and 5.1-times greater N fixation rates than roots of the tall phenotype under aerobic and anaerobic conditions, respectively (Figure 3). Conversely, sediment from the tall *S. alterniflora* phenotype presented greater potential nitrification rates between one and two orders of magnitude than all other studied compartments (Figure 3). In the short *S. alterniflora*, potential nitrification rates were an order of magnitude greater in root tissue compared to bulk and rhizospheric sediment (Figure 3).

**Figure 3.**
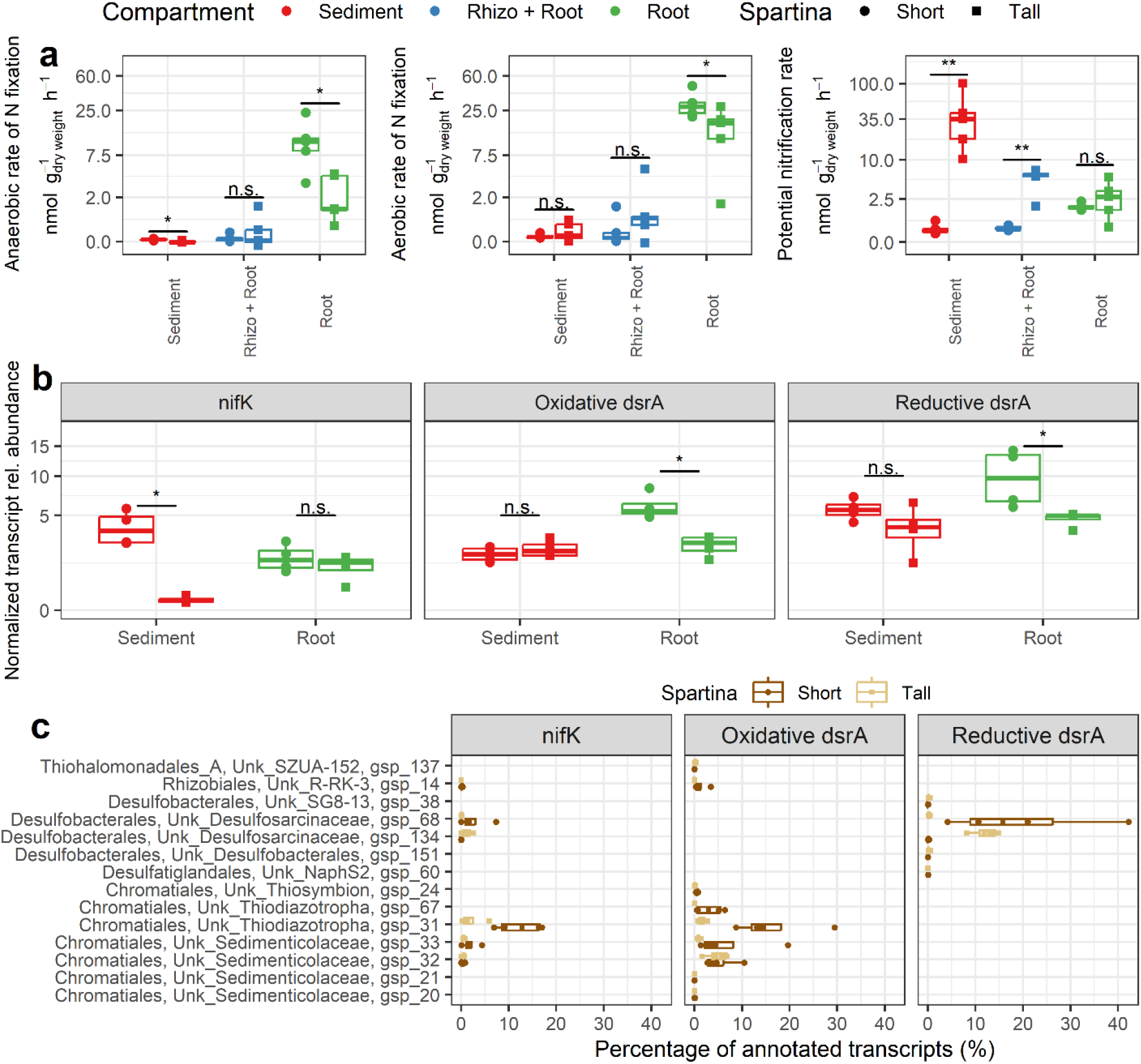
Drivers and metagenome assembled genomes (MAGs) associated with rates of N fixation and nitrification in the salt marsh environment. Rates of nitrification and N fixation (under anoxic and oxic conditions) per microbiome compartment and *Spartina alterniflora* phenotype (a). Normalized transcript relative abundance of the nitrogenase gene (nifK), and the reductive and oxidative dsrA types per microbiome compartment and *Spartina alterniflora* phenotype (b). Percentage from the total nitrogenase (nifK) and reductive and oxidative types of the dsrA transcripts mapping to the most active MAGs (c). Statistical significance based on non-parametric pairwise Mann-Whitney tests. n.s = p-value > 0.5, * p-value < 0.5, ** p-value < 0.01.

We used metagenomic and metatranscriptomic short reads to relate nitrogen fixation activity with multi-omics functional information. Metagenomic and metatranscriptomic reads were functionally annotated using the eggNOG database. We normalized the functional profile of each library to account for differences in sequencing effort and genome size by dividing the count matrix against the median abundance of 10 universal single-copy phylogenetic marker genes as in Salazar et al. (2019). Statistical difference between the normalized gene expression of the nitrogenase gene nifK and the oxidative and reductive dsrA types was calculated between *S. alterniflora* phenotypes across all compartments. Since gene expression within metabolic pathways was highly correlated, we used nifK and dsrA as marker genes (Supplementary Figure S3). Even though nitrogen fixation rates were the greatest in the root compartment of the short *S. alterniflora* phenotype, no statistical significance was found in the nitrogenase gene expression between the two *S. alterniflora* phenotypes (Figure 3). Conversely, the root compartment of the short *S. alterniflora* phenotype had the highest normalized transcript abundance of both reductive and oxidative dsrA gene types (Figure 3). To infer which members of the *S. alterniflora* root microbiome contributed the most to nitrogen fixation, as well as to test if nitrogen fixation was coupled to sulfur metabolism, we mapped all short read transcripts annotated as nifK and dsrA back to our binned gspp. Sulfur-oxidizing *Ca.* Thiodiazotropha gspp in the short phenotype of *S. alterniflora* contributed approximately 25% and 15% of all oxidative dsrA and nifK transcripts, respectively (Figure 3). In addition, the most active sulfur-oxidizing and sulfate-reducing gspp in roots of the short phenotype (gsp 31 and gsp 68) had a positive correlation between nitrogen fixation and sulfur oxidation/reduction gene expression, respectively (Supplementary Figure S4). In the tall phenotype, *Desulfosarcinaceae* gsp 134 contributed the most transcripts of the reductive dsrA gene. However, in general, we were not able to recover most of the nifK genes as part of the MAGs; and thus, transcripts, presumably due to issues with contig binning. Nevertheless, using short read analysis, we found that the majority of the nifK transcripts in the roots of the tall phenotype were assigned to bacteria from the *Desulfobacterota* phylum, while in the short phenotype most nifK transcripts were affiliated with bacteria from the *Gammaproteobacteria* class or the *Desulfobacterota* phylum (Supplementary Figure S5). Similarly, abundant transcripts of oxidative dsrA in the roots of the short phenotype were mostly affiliated with the *Gammaproteobacteria* class, while the reductive dsrA affiliated mainly with the *Desulfobacterales* and *Desulfovibrionales* orders in both phenotypes (Supplementary Figures S6 and S7).

### 2.2 Phylogeny, genetic potential and gene expression of microorganisms enriched in the *S. alterniflora* root microbiome

To assess the phylogenetic novelty and relation of *Sedimenticolaceae* sulfur-oxidizing MAGs from the present study in comparison to that of marine invertebrate chemosymbionts, we retrieved all publicly available *Sedimenticolaceae* genomes as reported by GTDB release R07-RS207, as well as all *Ca.* Thiodiazotropha spp. analyzed by Osvatic et al. (2023). A maximum-likelihood phylogenetic tree using a 400 universal marker genes database was performed in PhyloPhlAn. We found that *S. alterniflora* root symbionts assigned to the *Ca.* Thiodiazotropha genus and an unknown genus from the *Sedimenticolaceae* family formed a monophyletic clade with lucinid clam chemosymbionts (Figure 4). All recovered MAGs that were phylogenetically related to marine invertebrate chemosymbionts were highly abundant in the root compartment of the *S. alterniflora* short phenotype, where sulfidic conditions are found in the marsh environment (Figure 1). To conserve phylogenetic coherence, we propose that *S. alterniflora* root symbionts assigned to the *Sedimenticolaceae* family and that formed a monophyletic group with *Ca.* Thiodiazotropha spp. are also members of such genus (Figure 4). Out of the 7 recovered *Ca.* Thiodiazotropha gspp, 6 had genes for carbon fixation (RuBisCO), 5 had the complete or partial nifDKH nitrogenase genes for nitrogen fixation, and all of them harbor the complete or at least partial genes for dissimilatory sulfite reductase (oxidative dsrAB type), and sulfur oxidation by the SOX complex (soxABYXYZ). Furthermore, we calculated the gene expression profile of the most active *Ca.* Thiodiazotropha gspp (gsp 31 and gsp 33), finding that genes for sulfur oxidation through the oxidative dsrAB, soxABYXYZ complex, and cytochrome c oxidase cbb3-type genes, as well as genes for carbon and nitrogen fixation (RuBisCO and nifKDH) were among the most highly transcribed (Supplementary Tables S4 and S5). We also found that sulfate-reducing MAGs from the *Desulfosarcinaceae* family were enriched in the root compartment of *S. alterniflora* (Supplementary Figure S8). *Desulfosarcinaceae* gspp had contrasting preferences for root colonization, with gsp 68 mostly enriched in the roots of the short phenotype, gsp 134 preferentially enriched in the root of the tall phenotype, and gsp 80 equally abundant in both phenotypes (Supplementary Figure S8). Of the three *Desulfosarcinaceae* gspp, all contained the complete or partial genes for nitrogen fixation, and only gsp 80 did not have the reductive dsrAB gene. The most transcribed genes from both gsp 68 and gsp 134 were related to dissimilatory sulfate reduction (dsrAB and aprAB, Supplementary Tables S6 and S7).

**Figure 4.**
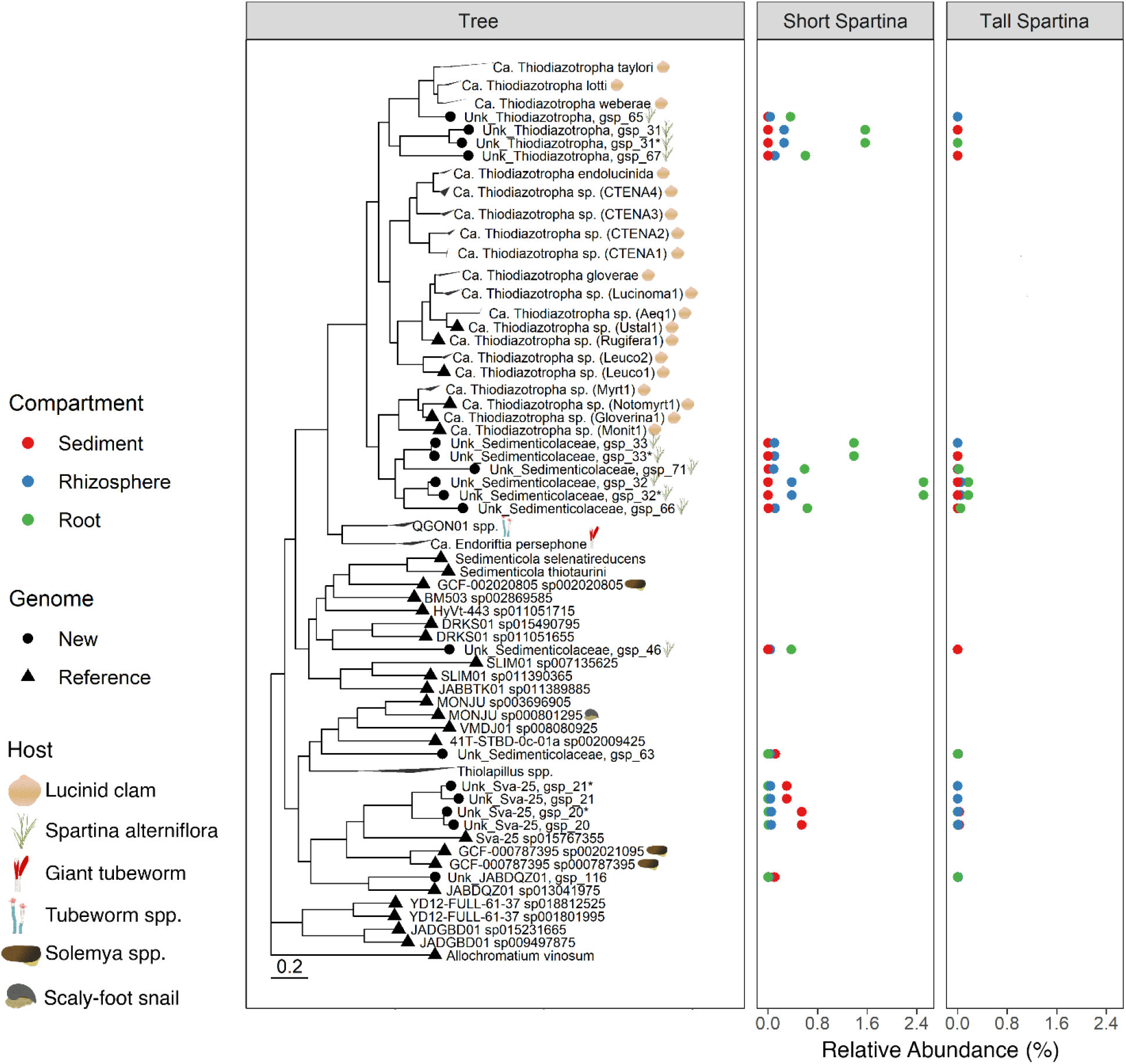
Phylogenetic tree of the Sedimenticolaceae family. Taxonomic annotation of the *Candidatus* Thiodiazotropha genus was based on Osvatic et al. (2023). The diagram next to the species name recognizes genomes recovered as symbionts of eukaryotic organisms. The average relative abundance of the metagenome assembled genomes (MAGs) from the present study is shown in the adjacent panels per microbiome compartment and *Spartina alterniflora* phenotype. Relative abundance was calculated at the DNA-level based on average coverage per position in metagenomic libraries. Purple sulfur bacteria *Allochromatium vinosum* was used as an outgroup for the phylogenetic tree.

### 2.3 Sediment and root microbiomes have contrasting functional profiles across a salt marsh primary productivity gradient

A decreasing trend in prokaryotic alpha divestity was observed from the sediment compartment to the root tissue in both *S. alterniflora* phenotypes (Figure 5). Similarly, quantification of the 16S rRNA gene by qPCR showed statistically lower prokaryotic abundance in the root tissue compared to the sediment and rhizosphere compartments (Figure 5). A stronger barrier for prokaryotic root colonization, evidenced by a steeper decrease in prokaryotic abundance in the rhizosphere-root interface, was observed in the less stressed tall *S. alterniflora* phenotype (Figure 5). Conversely, the root compartment of both *S. alterniflora* phenotypes presented a higher transcript copy number of the prokaryotic 16S rRNA than their sediment counterparts, indicating greater prokaryotic activity in the root zone (Figure 5).

**Figure 5.**
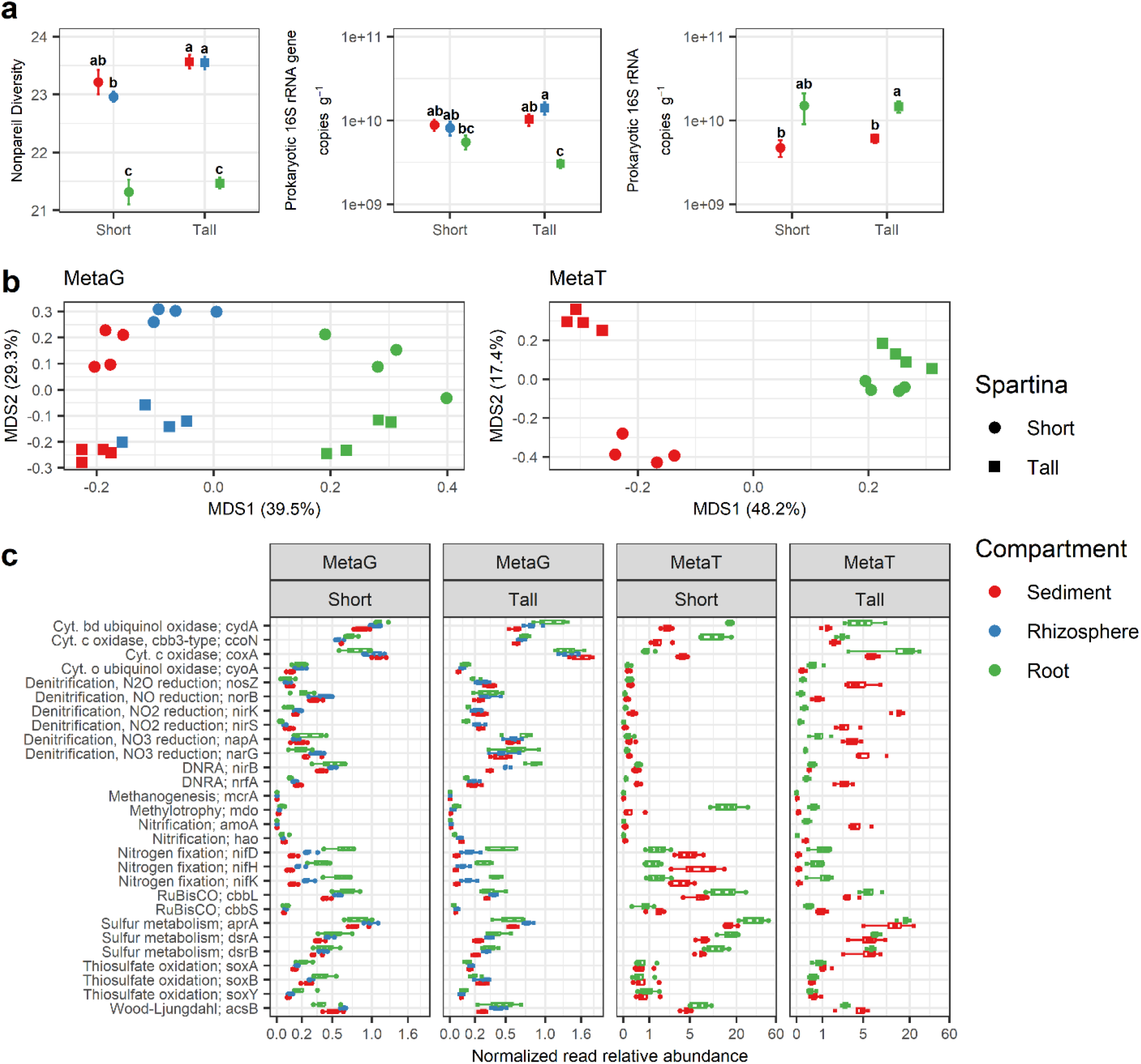
Prokaryotic abundance, functional diversity, and activity are determined by *Spartina alterniflora* phenotype and microbiome compartment. Average ± standard error of the metagenomic nonpareil diversity, 16S rRNA gene and transcript abundance as quantified by qPCR and RT-qPCR, respectively (n = 4) (a). Principal coordinate analysis (PCoA) ordination plot based on the Bray-Curtis dissimilatory index of functional profiles from KEGG orthology annotations (KO level) of metagenome and metatranscriptome libraries (b). Gene and transcript normalized relative abundance of prokaryotic terminal oxidases and selected enzymes of the carbon, nitrogen, and sulfur cycles (c). Different letter indicate statistical difference based on pairwise Mann-Whitney tests (p-value < 0.5).

PERMANOVA and principal coordinate analysis from metagenomic and metatranscriptomic functional profiles indicated that the environmental stress gradient significantly affects the microbiome’s potential and expressed functional repertoire (Figure 5, Supplementary Table S8). The community-level abundance and expression of terminal oxidase genes for aerobic respiration were specific to compartment and *S. alterniflora* phenotype. The relative abundance of genes for the canonical cytochrome c oxidase system (coxA) was greater in the tall *S. alterniflora* phenotype in both metagenomic and metatranscriptomic samples, while cbb3-type cytochrome c genes (ccoN) were mostly present and transcribed in the root tissue of the short *S. alterniflora* phenotype (Figure 5). Genes related to the redox cycling and turnover of nitrogen were more abundant and transcribed in the sediment of the tall *S. alterniflora* phenotype than all other assessed marsh compartments [i.e., nitrification, denitrification, and dissimilatory nitrate reduction to ammonium (DNRA) genes] (Figure 5). Conversely, genes for prokaryotic carbon fixation by RuBisCO and nitrogen fixation had the greatest relative abundance in the root compartment, especially in those from the short *S. alterniflora* phenotype (Figure 5). Nevertheless, relative gene expression for nitrogen fixation was greater in the sediment from the short phenotype and contributed mostly by cyanobacteria species (Supplementary Figure S5). Finally, greater redox cycling of sulfur in the root compartment, especially in the short *S. alterniflora,* was evidenced by the abundance and expression of sulfur dissimilatory metabolism genes (Figure 5).

### 2.4 Plant species from coastal marine ecosystems harbor unique root microbiomes

In order to understand how root microbiomes are assembled in coastal marine ecosystems compared to better-studied terrestrial plants, we used publicly available sequences to generate and curate a 16S rRNA gene amplicon database from root microbiomes comprising 2,911 amplicon samples from 56 plant species (complete dataset in Supplementary Table S9). We grouped amplicon samples into 4 broad ecosystem types: seabed, coastal wetlands, freshwater wetlands, and other terrestrial ecosystems. Species exchange based on the Bray-Curtis dissimilarity matrix was largely explained by ecosystem type, with coastal wetland and seabed ecosystems clustering together in an NMDS ordination (Figure 6). PERMANOVA analysis revealed that ecosystem type alone significantly explained 13.2% of the variation of the species exchange between analyzed samples (Supplementary Table S10). Furthermore, to explore two functional guilds that may explain the difference in microbiome assembly between ecosystem types, we assigned putative sulfate reduction and sulfur oxidation functions to our taxonomy table as performed by Rolando et al. (2022). We found that the root microbiomes of both seabed and coastal wetland ecosystems were highly enriched in putative sulfate-reducing and sulfur-oxidizing bacteria (Figure 6). Furthermore, we discovered that in both seabed and coastal wetland ecosystems, amplicons showing high sequence identity to known sulfur-oxidizing bacteria with the capability for nitrogen fixation (*Sedimenticolaceae* family: *Ca.* Thiodiazotropha genus) were highly abundant (Supplementary Figure S9). Conversely, amplicons showing high sequence identity to sulfate reducers from the *Desulfosarcinaceae* family, particularly those from the genus *Desulfatitalea* were highly enriched in the roots of coastal wetland plants (Supplementary Figure S9). The taxonomic identity of highly abundant ASVs in the root compartment of coastal marine plants coincide with diazotrophic MAGs retrieved from Sapelo Island, GA, such as sulfur oxidizers of the *Sedimenticolaceae* family, and sulfate reducers of the *Desulfosarcinaceae* family.

**Figure 6:**
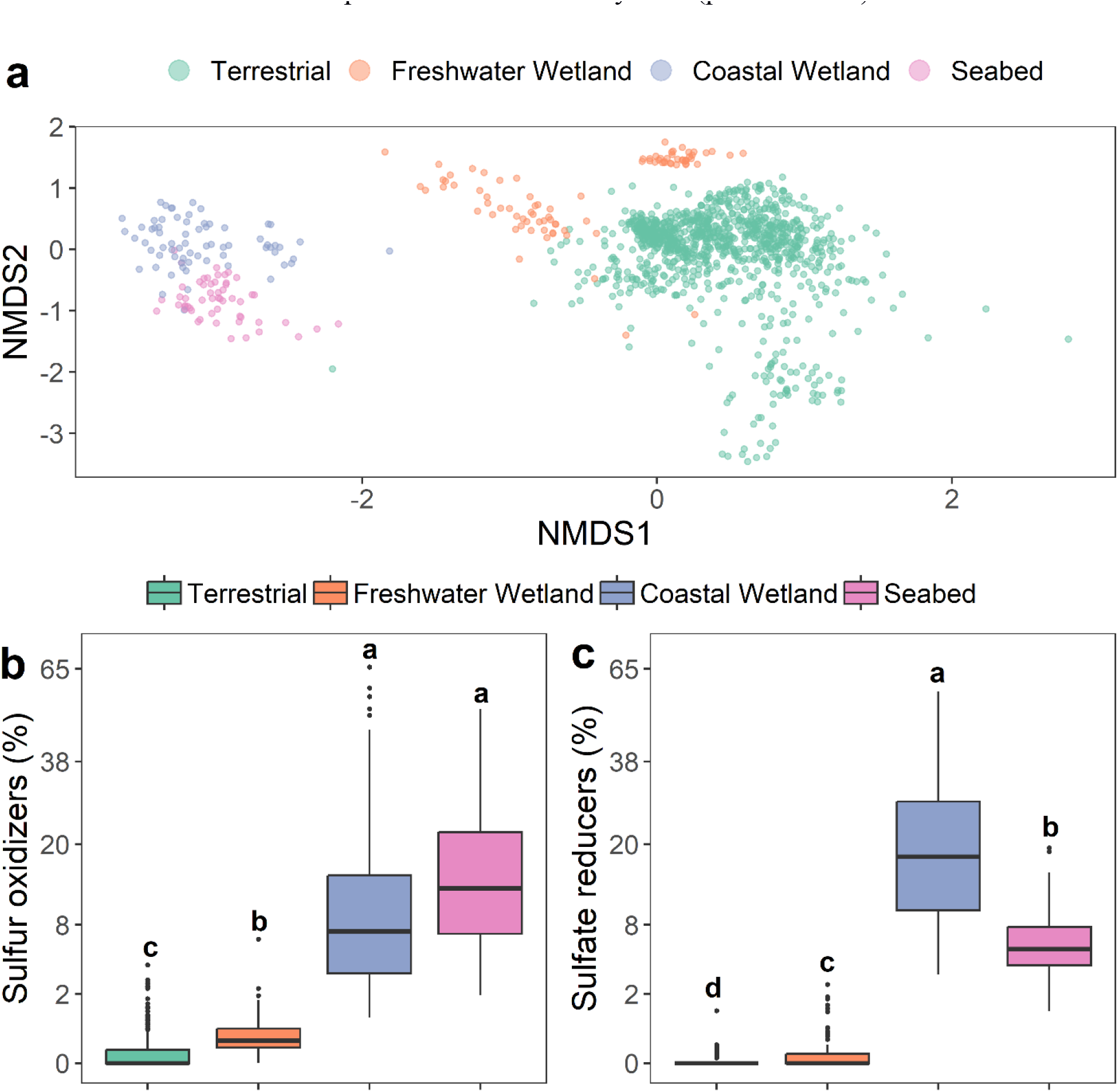
Roots from marine influenced ecosystems assemble a distinct microbial community enriched by bacteria with known to conserve energy from sulfur metabolism. Non-metric multidimensional scaling (NMDS) ordination plot based on the Bray-Curtis dissimilatory index of root-associated prokaryotic communities at the genus level, colored by ecosystem type (a). Relative abundance of putative sulfur-oxidizing (b) and sulfate-reducing root bacteria (c) by ecosystem type. Prokaryotic communities were characterized by analyzing an SSU rRNA gene amplicon dataset of 1182 samples assessing roots from 56 plant species. Different letter indicate statistical difference based on pairwise Mann-Whitney tests (p-value < 0.5).

## 3. Discussion

### 3.1 Nitrogen fixation is stimulated by rapid sulfur cycling in the roots of *S. alterniflora*

Nitrogen fixation in belowground coastal vegetated ecosystems has been mainly associated with heterotrophy, and particularly with sulfate reduction (Gandy and Yoch, 1988; McGlathery et al., 1998; Herbert, 1999; Nielsen et al., 2001; Welsh et al., 1996). The linkage of sulfate reduction to nitrogen fixation was proposed based on studies quantifying rates of diazotrophy with and without sulfate reduction inhibition by molybdate. Here, we show for the first time that highly-abundant sulfur-oxidizing bacteria in the roots of *S. alterniflora* also encode and highly express genes for nitrogen fixation (Figure 2, Figure 3). Furthermore, the MAG with the highest nitrogenase gene expression in our study was a novel sulfur-oxidizing bacterium from the *Ca.* Thiodiazotropha genus that also showed high expression of sulfur oxidation genes (Figure 3, Supplementary Figure S4). Our results suggest that when sulfate reduction was inhibited in previous studies, it disrupted the redox cycling of sulfur by impeding the flow of reduced sulfur compounds that serve as an energy source for sulfur-oxidizing bacteria. Thus, most likely previous inhibition studies not only blocked nitrogen fixation from sulfate-reducing microorganism, but also significantly reduced nitrogen fixation coupled to sulfur oxidation. Unlike sulfate reduction, there is no reliable method for measuring sulfide oxidation rates. Thus, the abundance and expression of sulfur-oxidation genes is one of the few ways to directly link this otherwise unmeasurable process with nitrogen fixation. We propose that both dissimilatory sulfur reduction and oxidation, and more importantly, the rapid redox cycling of sulfur, stimulate nitrogen fixation in the root environment of *S. alterniflora*. Since coastal wetland plants and seagrasses assemble similar root microbiomes (Figure 6), and macrophyte activity boosts both nitrogen fixation and the cycling of sulfur, this may be a common phenomenon in the root zone of many coastal marine plants (Whiting et al., 1986; McGlathery et al., 1998; Welsh et al., 1996).

Further evidence supporting the significance of the rapid cycling of sulfur coupled to nitrogen fixation is that members of the *Gammaproteobacteria* class and *Desulfobacterota* phylum contributed most of the nitrogenase and oxidative/reductive dsrAB transcripts in the root compartment of *S. alterniflora*. Similar results have been obtained from DNA and RNA amplicon studies in seagrass and other coastal wetland plant species, where nitrogen fixation genes and their expression have been affiliated with microorganism closely related to sulfate-reducing and sulfur-oxidizing bacteria (Thomas et al., 2014; Crump, et al. 2018; Kolton et al., 2020).

### 3.2 The root zone of coastal marine ecosystems is a hotspot of sulfur redox cycling

In most coastal marine ecosystems, water saturated sediments are depleted in oxygen within the first millimeters depth (Jørgensen et al., 2019). Because coastal marine ecosystems are bathed in seawater containing high sulfate concentrations (28 mM), sediment sulfate-reducing microorganisms often perform the terminal step of organic matter decomposition (Jørgensen et al., 2019). Unlike aerobic respiration, energy flow during sulfate reduction is decoupled from the carbon cycle, with most of the free energy conserved in reduced sulfur compounds (Howarth, 1984). The chemically stored energy is subsequently released by biotic and abiotic oxidation reactions at marsh oxic-anoxic interfaces coupled to the reduction of potent electron acceptors (oxygen, nitrate, or oxidized metals) (Howarth, 1984). Surface sediments of vegetated coastal marine ecosystems experience rapid re-oxidation of most reduced sulfur compounds. This process is highly concentrated in the microaerophilic root zone, which serves as a hotspot for the reaction (Holmer et al., 2002; Kristensen and Alongi, 2006; Koop-Jakobsen et al., 2018). Our results showing that the root compartment of coastal marine ecosystems is highly enriched in microorganisms with the capability for both sulfur reduction and oxidation provides further evidence for rapid sulfur cycling in the root zone (Figure 6). Furthermore, to the best of our knowledge, this is the first study to employ genome reconstruction from metagenomic sequencing to verify the function of uncultivated sulfate-reducing and sulfur-oxidizing microorganisms living in the roots of coastal wetland plants. Our findings also reveal that genes that catalyze oxidative and reductive reactions in the sulfur cycle are highly transcribed in the root environment, particularly in the stressed *S. alterniflora* phenotype that thrives in a sulfidic environment (up to 1.5 mM sulfide concentration in our studied transects; Rolando et al. 2022). This is consistent with previous studies of salt marsh and seagrass ecosystems, in which genes of sulfur-oxidation pathways were shown to be highly expressed in the root compartment of *S. alterniflora* and *Zostera* seagrass spp., respectively (Thomas et al. 2014, Crump et al., 2018). Thus, we propose that in contrast to terrestrial ecosystems, coastal marine plants rely on the rapid cycling of sulfur in their root zone for the breakdown of organic matter and recycling of nutrients through sulfate reduction along with the re-oxidation of terminal electron acceptors by sulfur oxidation. Furthermore, we propose that rapid rates of both oxidative and reductive sulfur reactions represent a key mechanism to sustain high rates of energetically-expensive nitrogen fixation in the root zone of *S. alterniflora*.

### 3.3 Phylogenetic and ecological relationship between marine invertebrates and *S. alterniflora* chemosymbionts

Endosymbionts from the *Ca.* Thiodiazotropha genus were initially discovered living in symbiosis with lucinid clams, where they fix carbon and serve as a source of carbon and nitrogen to the animal host using energy gained from the oxidation of reduced forms of sulfur (Petersen et al., 2017). Recent studies using fluorescence in situ hybridization (FISH) microscopy and SSU rRNA gene metabarcoding have shown that *Ca.* Thiodiazotropha bacteria also inhabit the roots of a diverse array of seagrass species along with that of the coastal cordgrass *S. alterniflora* (Martin et al., 2020a; Rolando et al., 2022). Here, we show that close relatives of diazotrophic sulfur chemosymbionts associated with lucinid clams were highly abundant and active in the roots of *S. alterniflora*. To the best of our knowledge, this is the first study reporting genomes and gene expression profiles from *Ca.* Thiodiazotropha retrieved from macrophyte root samples. We show that our MAGs formed a monophyletic clade with those from previously described marine invertebrate animals (Figure 4). Similar to what has been reported in their symbiosis with lucinid clams, the *S. alterniflora* symbionts highly expressed genes for sulfur oxidation, and carbon and nitrogen fixation (Supplementary Table S4, Supplementary Table S5; Petersen et al., 2017; Lim et al., 2019a). Studies of lucinid clams have shown that the *Ca.* Thiodiazotropha symbionts are horizontally transmitted (Petersen and Yuen, 2021). Furthermore, invertebrate colonization by *Ca.* Thiodiazotropha symbionts is not restricted by either the host or symbiont species (Lim et al., 2019a). The flexibility of this symbiotic relationship could explain the evolution of *Ca.* Thiodiazotropha symbionts colonizing macrophyte roots. Our results show that all *Ca.* Thiodiazotropha symbionts of *S. alterniflora* are distinct species from those previously found in lucinid clam hosts (ANI < 82% for all pairwise comparisons). However, additional metagenomic sequencing of both macrophyte plants and lucinid clams is needed to better interrogate if chemosymbionts populations are restricted by host biology at the kingdom level (i.e., to either plant or animal hosts). In contrast to what is observed in lucinid clams, where a single bacterial species dominates the gill microbiome; in *S. alterniflora* and seagrass species, *Ca.* Thiodiazotropha spp. do not completely outcompete other microbial species in the root compartment (Lim et al., 2019a, 2019b). However, the symbionts comprise a large proportion of the microbial community, and are amongst the most active species (Rolando et al., 2022; Supplementary Figure S9).

### 3.4 Sulfur chemosymbiosis as an adaptation to anoxia and sulfide toxicity in *S. alterniflora* plants

We propose that the symbiosis between *S. alterniflora* and sulfur-oxidizing bacteria represents a key adaptation supporting the resilience of coastal salt marsh ecosystems to environmental perturbations. For instance, intertidal wetland ecosystems are vulnerable to climate change because they are located in a narrow elevation range determined by tidal amplitude (Morris et al., 2021). The Intergovernmental Panel on Climate Change (IPCC) has projected an increase in sea level between 0.38 m and 0.77 m by 2100 (IPCC-AR6, Fox-Kemper et al., 2021). Although coastal wetland ecosystems are dynamic and adapt to sea level rise by increasing sediment accretion rates, ecosystem models predict that a large area of present-day marsh will drown because of accelerated sea level rise (Kirwan et al., 2016). Increased hydroperiods will impact the redox balance of vegetated sediments, imposing more severe anoxia and physiological stress from sulfide toxicity to wetland plants. Under this scenario, a symbiosis of coastal vegetated plants with sulfur-oxidizing bacteria could alleviate sulfide stress while at the same time coupling it to carbon and nitrogen fixation for potential plant uptake; however, the mechanism for carbon and/or nitrogen transfer between sulfur-oxidizing symbionts and the host plants still remains elusive and requires further research. Similar to Thomas et al. (2014), we showed the transcription of sulfur oxidation genes was greater in the stressed *S. alterniflora* short phenotype. In seagrass ecosystems, Martin et al. (2020b) also found that the seagrass root microbiome was more enriched in *Ca.* Thiodiazotropha spp. under stress conditions. Thus, we suggest that the *S. alterniflora* – *Ca.* Thiodiazotropha symbiosis is an adaptive interaction to anoxic soil conditions, whereby the host plant responds to stress from elevated dissolved sulfide concentrations. Further, we show that the short *S. alterniflora* phenotype, which harbors a greater nitrogenase transcript abundance of *Ca.* Thiodiazotropha sulfur chemosymbionts, also had the greatest rates of N fixation of all assessed marsh compartments. Our results support the SGH in which greater stress strengthens the relationship between sulfur oxidizing bacteria and macrophyte plants. Stressed plants benefit from the symbiotic relationship through reduced sulfide toxicity and the coupling of sulfide oxidation to nitrogen fixation, with nitrogen likely transferred to the plant host. Conversely, the tall *S. alterniflora* phenotype showed a faster internal cycling of nitrogen, which is consistent with previous studies showing greater rates of nitrogen mineralization in this zone of the ecosystem (Figure 5, Rolando et al., 2022).

Intriguingly, we also found high transcript expression of RuBisCO from *Ca.* Thiodiazotropha gspp in the root compartment of the stressed short phenotype of *S. alterniflora* (Supplementary Table S5). When assessing the natural abundance of ^13^C (δ^13^C) in the roots of the short *S. alterniflora*, we found that it was significantly depleted compared to the δ^13^C from the tall phenotype (Supplementary Figure S10). As a C4 plant, *S. alterniflora* first fixes carbon by the action of PEP carboxylase, which has lower discrimination against ^13^C than RuBisCO (Farquhar, 1983). Thus, we could speculate that the higher ^13^C discrimination in the root of the short *S. alterniflora* phenotype is a consequence of sulfur chemoautotrophic carbon fixation by RuBisCO. Consistent with our hypothesis, a previous study by Hwang and Morris (1992) showed that the roots of *S. alterniflora* fix carbon through an undeciphered dark reaction. Further, unlike other plant species, *S. alterniflora* assimilates sulfur as sulfide (Carlson and Forrest, 1982), and under mild sulfidic conditions, sulfur oxidation in *S. alterniflora* roots is mostly biological and was previously hypothesized to be coupled with energetic gain (Lee et al., 1999). Interestingly, low doses of sulfide (< 1 mM) have been shown to stimulate the productivity of *S. alterniflora* (Mendelssohn and Morris, 2000). However, since other plant physiological processes could also explain differences in δ^13^C, such as differing rates of dark respiration, and leakiness from the bundle sheath cells to the mesophyll cell (Farquhar, 1983), further investigations are required to quantify the relevance of sulfur chemolithoautotrophy in *S. alterniflora* root biomass production.

## 4. Materials and methods

### 4.1 Study site, field sampling and sample processing

The field fraction of this study was carried out in the Georgia Coastal Ecosystem - Long Term Ecological Research (GCE-LTER) site 6, located on Sapelo Island, GA (Lat: 31.389**°** N, Long: 81.277**°** W). Four ∼100 m transects along the tall to short *Spartina alterniflora* gradient were studied in July 2018, 2019, and 2020 (Figure 1, further site description in Rolando et al., 2022). A combination of multi-omics approaches and biogeochemical rate measurements were performed across different salt marsh compartments and phenotypes of *S. alterniflora*.

The microbiome of *S. alterniflora* was analyzed using shotgun metagenomics from three compartments: sediment, rhizosphere, and root. The tall and short *S. alterniflora* extremes of the four studied transects were sampled (n: 4 transects * 2 *S. alterniflora* phenotypes * 3 compartments = 24). Samples from transects 1 and 2 were collected in July 2018, while those from transects 3 and 4 were collected in July 2019. All sampling was performed at the 0-5 cm depth profile. Root-associated samples were washed two times with creek water in the field to remove coarse chunks of sediment attached to the plant. All samples were immediately flash-frozen in an ethanol and dry ice bath, and stored at −80°C until DNA extraction.

Samples for metatranscriptomic analysis and biogeochemical rate measurements were collected in July 2020 only from transect 4 to reduce plant disturbance time before flash freezing and incubations, respectively. Five independent plants at least 3 meters apart were sampled with a shovel at the two *S. alterniflora* biomass extremes. The entire plant, including the undisturbed root system with sediment attached, was sampled up to at least 20 cm depth, transferred to a 5 gallons bucket, and immediately transported to the field lab. For each plant, a paired sample of sediment was collected. In the lab, roots were washed with creek water two times. After the first wash, roots with sediment attached were collected and defined as the rhizosphere + root compartment. Live, sediment-free roots were sampled after the second wash and defined as the root compartment. Top 5 cm from each sample were homogenized and used for total RNA extractions and measurements of nitrogen fixation and potential nitrification rates. Root samples for RNA extractions were washed in an epiphyte removal buffer, as in Simmons et al. (2018). Sediment and root samples for RNA analysis were immediately flash-frozen in an ethanol dry ice bath, and stored at −80 °C until extraction. Due to limited sample amount, we did not prepare rhizosphere libraries for metatranscriptomic analysis.

### 4.2 Nitrogen fixation and potential nitrification rates

Rates of N fixation were calculated in ^15^N_2_ incubations in 14 ml serum vials as in Leppänen et al. (2013). Rates were measured for all samples under both oxic and anoxic conditions. About 2 g, and 1 g of wet weight was used for tracer gas incubations of sediment and root samples, respectively. An additional subsample of 2 g was immediately oven-dried at 60 °C for 72 hours to be used as a pre-incubation control. In all vials containing root samples, 4 ml of autoclaved and 0.22 um filter-sterilized artificial seawater was added to avoid tissue desiccation. No artificial seawater was added to sediment and rhizosphere samples since they were already water-saturated. In anoxic incubations, vials were flushed with N_2_ gas for 5 minutes. After sealing all incubation vials, 2.8 ml of gas was removed and immediately replaced with 2.8 ml of ^15^N_2_ (98% enriched, Cambridge Isotope Laboratories Inc, USA). Vials were overpressured by adding 0.3 ml of air or N_2_ gas in oxic or anoxic incubations, respectively. The incubations were carried out in the dark and at room temperature for 24 hours. At the end of the incubation, samples were oven dried at 60 °C for 72 hours, and ground using a PowerGen high throughput homogenizer (Fisherbrand, Pittsburgh, PA). Ground samples were sent to the University of Georgia Center for Applied Isotope Studies (https://cais.uga.edu/) for carbon and nitrogen elemental analysis and ^13^C and ^15^N stable isotope analysis. Elemental analysis was performed by the micro-Dumas method, while isotopic analysis by isotope ratio mass spectrometry. Rates of ^15^N incorporation were calculated per dry weight basis (DW) as in Leppänen et al. (2013):

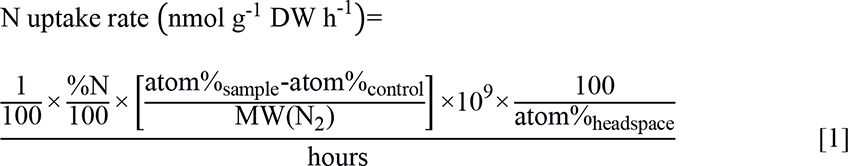

Where %N is the N percent concentration of the oven-dried sample, and MW(N_2_) is the molecular weight of N_2_ (28.013446).

Potential nitrification rates were measured as detailed by Dollhopf et al. (2005) in 5 replicates for each tall and short *S. alterniflora* compartments: sediment, rhizosphere + root, and root. Homogenized samples (5 g, 5 g, and 2 g for sediment, rhizosphere + root, and root compartment, respectively) were placed in 250-ml Erlenmeyer flasks filled with 50 ml sterile artificial seawater. Sodium chlorate and ammonium sulfate were amended to all flasks to final concentrations of 10 mM and 500 µM, respectively. Flasks were incubated in the dark at room temperature and orbital shaking (115 rpm) for 24 hours. Subsamples for NO_2_ analysis were taken at 0, 4, 8, 12, 18, and 24 hours. Timepoint samples were filtered with 0.2 µm syringe filters, and stored at −20 °C until analysis. Filtered NO_2_ concentration was measured by spectrophotometry (García-Robledo et al., 2014). Potential nitrification rates (N nmol g^-1^_DW_ h^-1^) were calculated as the slope of the linear regression of NO_2_ concentration over time.

### 4.3 Nucleic acid extractions and multi-omics library preparation

For DNA extractions, compartment separation of the rhizosphere and root microbiomes was performed by sonication in an epiphyte removal buffer, as detailed in Simmons et al. (2018). Extracellular dissolved or sediment-adsorbed DNA was removed from bulk sediment samples according to the Lever et al. (2015) procedure. DNA extractions from all samples were performed using the DNeasy PowerSoil kit (Qiagen, Valencia, CA) following the manufacturer’s instructions. Shotgun metagenome sequencing was performed on an Illumina NovaSeq 6000 S4 2x150 Illumina flow cell at the Georgia Tech Sequencing Core (Atlanta, GA).

RNA from 4 biological replicates of both the sediment and root compartments from the tall and short phenotypes of *S. alterniflora* was extracted using the ZymoBIOMICS RNA Miniprep (Zymo Research Corp) kit according to the manufacturer’s protocol (n: 4 replicates * 2 *S. alterniflora* phenotypes * 2 compartments = 16 samples). Rigorous DNA digestion was done with the TURBO DNase kit (Invitrogen), and eukaryotic mRNA was removed by binding and discarding the eukaryotic mRNA polyA region to oligo d(T)25 magnetic beads (England Biolabs). Finally, rRNA from both plant and prokaryotic organisms was depleted using the QIAseq FastSelect -rRNA Plant, and -5S/16S/23S kits, respectively (Qiagen, Valencia, CA). Metatranscriptomic libraries were sequenced in two lanes of Illumina’s NovaSeq 6000 System flow cell utilizing the NovaSeq Xp workflow (SE 120bp) at the Georgia Tech Sequencing Core (Atlanta, GA). We retrieved 203, and 415 Gpb of metagenomic and metatranscriptomic raw sequences, respectively; with a median sequencing effort of 7.8, and 25.3 Gbp per metagenomic and metatranscriptomic library (Supplementary Table S1). After quality control and *in silico* removal of host and rRNA reads, median sequencing effort decreased to 5.3 and 9 Gbp per metagenomic and metatranscriptomic library, respectively (Supplementary Table S1).

### 4.4 Gene and transcript quantification of the prokaryotic 16S rRNA gene

Prokaryotic abundance and a proxy of prokaryotic activity were measured by quantitative polymerase chain reaction (qPCR) and Reverse Transcription-qPCR (RT-qPCR) of the SSU rRNA gene, respectively. All samples used for metagenome and metatranscriptome analysis were quantified by qPCR and RT-qPCR, respectively. Samples were analyzed in triplicate using the StepOnePlus platform (Applied Biosystems, Foster City, CA, USA) and PowerUp SYBR Green Master Mix (Applied Biosystems, Foster City, CA, USA). Reactions were performed in a final volume of 20μl using the standard primer set for the prokaryotic SSU rRNA gene: 515F (5′-GTGCCAGCMGCCGCGGTAA′) and 806R (5′-GGACTACHVGGGTWTCTAAT′) (Caporaso et al., 2011, Rolando et al., 2022). To avoid plant plastid and mitochondrial DNA/cDNA amplification from root samples, peptide nucleic acid PCR blockers were added to all qPCR and RT-qPCR reactions at a concentration of 0.75 μM (Lundberg et al., 2013). Standard calibration curves were performed using a 10-fold serial dilution (10^3^ to 10^8^ molecules) of standard pGEM-T Easy plasmids (Promega, Madison, WI, USA) containing target sequences from *Escherichia coli* K12. Melting curve analyses was used to check for PCR specificity. Prokaryotic gene and transcript abundance of the SSU rRNA gene were calculated as gene and transcript copy number g^−1^ of fresh weight, respectively.

### 4.5 Metagenomic and metatranscriptomic quality control

Metagenomic and metatranscriptomic raw reads were quality trimmed (quality phred score < 20), and filtered for Illumina artifacts, PhiX, duplicates, optical duplicates, homopolymers, and heteropolymers using JGI’s BBTools toolkit (Bushnell, 2014). Reads shorter than 75 bp were removed, and the quality of both metagenomic and metatranscriptomic libraries was assessed with FastQC (Andrews, 2010). Remaining reads were mapped against the only publicly available *S. alterniflora* genome (NCBI BioProject: PRJNA479677) with bowtie2 (Langmead and Salzberg, 2012), followed by removal of short reads that aligned to the *S. alterniflora* genome using samtools (parameters: view -u -f 12 -F256, Li et al., 2009). In addition, rRNA reads from metatranscriptomic samples were removed using sortMeRNA v.4.3.4 (Kopylova et al., 2012). Finally, DNA contamination in metatranscriptomic samples was assessed as indicated by Johnston et al. (2019). Short-read transcripts were mapped to all assembled contigs, and strand-specificity (consistency in sense/antisense orientation) was calculated for all genes with more than 100 hits. All assessed samples had a greater than 95% average strand-specificity, thus considered to be free of DNA contamination (Supplementary Table S1). Filtered, quality-trimmed, and host-free reads were utilized for subsequent analyses.

### 4.6 Metagenomic and metatranscriptomic short reads functional analysis and nonpareil diversity

Short reads from all metagenomic and metatranscriptomic samples were aligned against the eggNOG protein database (release 5.0.2) using eggnog-mapper v.2.1.9 (Cantalapiedra et al., 2021). The DIAMOND tabular outputs were filtered by retrieving only the best hit based on bitscore. Hits with less than 30% identity, less than 30% match of the read length, or that did not match a prokaryotic domain were removed from the analysis. Gene profiles for each metagenomic and metatranscriptomic library were constructed by counting hits of predicted KEGG orthology. Due to differences in sequencing effort and genome size between multi-omics libraries, the count matrixes were normalized by dividing them by their median count of 10 universal single-copy phylogenetic marker genes (K06942, K01889, K01887, K01875, K01883, K01869, K01873, K01409, K03106, and K03110) as in Salazar et al. (2019). Functional gene and transcript profiles were analyzed by an principal coordinate analysis utilizing the Bray-Curtis dissimilarity distance. Further, for both metagenomic and metatranscriptomic profiles, multivariate variation of the Bray-Curtis dissimilarity matrix was partitioned to compartment (sediment, rhizosphere, and root) and *S. alterniflora* phenotype, based on a permutational multivariate analysis of variance (PERMANOVA) with 999 permutations performed in vegan v. 2.5.7 (Oksanen et al., 2013). The normalized relative abundance of selected terminal oxidases and genes/transcripts from the carbon, nitrogen, and sulfur cycles were assessed by compartment and *S. alterniflora* phenotype.

After removing reads annotated as Eukaryotic by eggnog-mapper, we calculated nonpareil diversity from all metagenomic samples using nonpareil v. 3.401 (Rodriguez-R et al., 2018b). Nonpareil diversity is a metric estimated from the redundancy of whole genome sequencing reads, and has been shown to be closely related to classic metrics of microbial alpha diversity like the Shannon index.

### 4.7 Recovery of metagenome assembled genomes (MAGs)

The following binning approach was motivated by a recently proposed method of iteratively subtracting reads mapping to MAGs, re-assembling, and re-binning metagenomic libraries to increase the number of recovered genomes (Rodriguez-R et al., 2020). An initial assembly using idba-ud v1.1.3 with pre-error-correction for highly uneven sequencing depth was performed for all individual metagenomic libraries (default parameters), as well as co-assemblies grouping libraries from the same *S. alterniflora* phenotype and compartment (--mink 40 --maxk 120 --step 20 --min_contig 300) (Peng et al., 2012). The resulting contigs from both individual and co-assemblies were binned using three different algorithms with default options: MaxBin v.2.2.7, MetaBAT v.2.15, and CONCOCT v1.1.0 (Alneberg et al., 2014; Wu et al., 2016; Kang et al., 2019). Recovered bins were dereplicated with DAS Tool v1.1.2 (Sieber et al., 2018), and the output was refined for putative contamination with MAGPurify v2.1.2 (Nayfach et al., 2019). Completeness, contamination, and quality score (Completeness – 5*Contamination) were calculated with MiGA v0.7 (Rodriguez-R et al., 2018a). Recovered MAGs with a quality score less than 50 were considered low quality and discarded. A second round of assembly and binning was performed after discarding short reads that mapped against high-quality (HQ) MAGs. Metagenomic libraries were mapped against HQ MAGs using bowtie2 with default parameters. Paired reads that mapped against HQ MAGs were discarded using samtools (parameters: view -F 2, Li et al., 2009). Filtered metagenomic libraries were co-assembled once again using idba-ud v1.1.3 with pre-error-correction for highly uneven sequencing depth (parameters: --mink 40 -- maxk 120 --step 20 --min_contig 300) in four groups: i) tall *S. alterniflora* root, ii) short *S. alterniflora* root, iii) tall *S. alterniflora* sediment and rhizosphere, and iv) short *S. alterniflora* sediment and rhizosphere. Binning, MAGs’ refinement, and quality control were performed as explained for the first iteration. Finally, HQ MAGs from both iterations were grouped into genomospecies (gspp, singular gsp) by clustering MAGs with ANI > 95% using the MiGA v0.7 derep_wf workflow (Rodriguez-R et al., 2018a). MAGs with the highest quality score were selected as representatives of their gsp, and most downstream analyses were performed with them.

Genomospecies relative abundance was estimated for each metagenomic library as in Rodriguez-R et al. (2020). Sequencing depth was calculated per position using bowtie2 with default parameters (Langmead and Salzberg, 2012), and bedtools genomecov (parameters: -bga, Quinlan and Hall, 2010). Bedtools output was truncated to keep only the central 80% values, and the mean of all retained positions was calculated using BedGraph.tad.rb from the enveomics collection, a metric defined as TAD_80_ (truncated average sequencing depth) (Rodriguez-R and Konstantinidis, 2016). Relative abundance of each gsp was calculated by dividing TAD_80_ by genome equivalents estimated for each metagenomic library with MicrobeCensus (Nayfach and Pollard, 2015).

### 4.8 MAGs taxonomy, phylogenetic analysis, and gene functional annotation and expression

Taxonomic classification of all MAGs was performed by GTDB-Tk v2.1.0 using the reference database GTDB R07-RS207 (Chaumeil et al., 2022). A maximum likelihood phylogenetic tree of all gspp was constructed using a 400 universal marker genes database in PhyloPhlAn v3.0.58 (parameters: -d phylophlan, --msa mafft, --trim trimal, --map_dna diamond, --map_aa diamond, - -tree1 iqtree, --tree2 raxml, --diversity high, --fast, Asnicar et al., 2020). The phylogenetic tree was decorated and visualized in ggtree v2.0.4 (Yu et al., 2018).

Protein-encoding genes from all binned MAGs were predicted with Prodigal v.2.6.3 using default parameters (Hyatt et al., 2010), and resulting amino acid sequences aligned against the eggNOG database (release 5.0.2) using eggnog-mapper v2.1.9 (Cantalapiedra et al., 2021). The DIAMOND tabular output was filtered by retrieving only the best hit based on bitscore, and removing hits with less than 30% identity and/or less than 50% match length.

Metagenomic and metatranscriptomic short reads were mapped against functionally annotated ORFs of all gspp using megablast v.2.10.1. Only hits with greater than 95% percent identity and 90% read alignment were retained. Percent contribution of genes and transcripts to the total microbial community was assessed by dividing the number of hits from each gsp against the total number of functionally annotated short reads (from section 4.7). When assessing the transcript expression profile of a specific gsp, the number of mapped hits per annotated transcript was normalized by dividing it by gene length (bp) and the median abundance of 10 universal single-copy phylogenetic marker genes of the prokaryotic community (K06942, K01889, K01887, K01875, K01883, K01869, K01873, K01409, K03106, and K03110).

Finally, since the phylogeny of the dsrAB gene allows to discriminate between the oxidative and reductive dsrAB types, we aligned all recovered genes from our binned gspp to a reference alignment (Müller et al., 2015) with Clustal Omega v.1.2.4 (Sievers et al., 2011), and built an approximately-maximum-likelihood phylogenetic tree with FastTree v.2.1.11 (Price et al., 2010). Gene type was inferred based on placement in the phylogenetic tree.

### 4.9 Phylogenetic reconstruction of the *Sedimenticolaceae* family

Publicly available genomes from all species from the *Sedimenticolaceae* family, according to GTDB R07-RS207, and all *Candidatus* Thiodiazotropha genomes from the Osvatic et al. (2023) study were retrieved from NCBI. A full list and characteristics of genomes used for this analysis is found in Supplementary Table S11. A phylogenetic tree of all retrieved genomes and binned genomes from this study was constructed in PhyloPhlAn v3.0.58 using a 400 universal marker genes database (parameters: -d phylophlan, --msa mafft, --trim trimal, --map_dna diamond, -- map_aa diamond, --tree1 fasttree, --tree2 raxml, --diversity low, --accurate, Asnicar et al., 2020). The phylogenetic tree was decorated and visualized using ggtree v2.0.4 (Yu et al., 2018).

### 4.10 Analysis of root microbiomes from contrasting ecosystems

Publicly available 16S rRNA gene amplicon datasets, generated from next-generation sequencing, were used to characterize the community assembly of root microbiomes from coastal marine ecosystems. Studies were selected based on google scholar queries using a combination of the following keywords: “salt marsh”, “coastal”, “wetland”, “mangrove”, “seagrass”, “seabed”, “crop”, “bog”, “plant”, “root”, “endosphere”, “microbial community”, “microbiome”, “amplicon”, “next-generation sequencing”, “16S rRNA”, and “SSU rRNA”, as well as based on the authors prior knowledge. Only studies that collected environmental samples were included (i.e., no greenhouse or plants grown on potting media were included). When available, paired soil/sediment and rhizosphere samples were also retrieved. Twenty-two studies that met our requirements were selected, collecting a total of 2911 amplicon samples, with 1182 of them being from the root compartment across 56 different plant species. Selected plants were categorized into 4 different ecosystem types: seabed, coastal wetlands, freshwater wetlands, and other terrestrial ecosystems. A complete list of selected amplicon samples with accompanying metadata is available in Supplementary Table S9.

Cutadapt v.3.7 was used to detect and remove primer sequences from the datasets (Martin, 2011). Primer-free sequences were quality filtered using DADA2’s filterAndTrim function [options: truncLen=c(175,150), maxN=0, maxEE=c(2,2), truncQ=10, rm.phix=TRUE] (Callahan et al., 2016). Trimmed reads were randomly subsampled to a maximum of 20,000 reads using seqtk v.1.3 (Li, 2012) and used as input for dada2 amplicon sequence variant (ASV) calling (Callahan et al., 2016). Chimeras were removed using the removeBimeraDenovo function from the DADA2 package. Taxonomy was assigned to ASVs utilizing the Ribosomal Database Project (RDP) Naive Bayesian Classifier (Wang et al., 2007) against the SILVA SSU rRNA reference alignment [Release 138, (Quast et al., 2012)]. Sequences classified as chloroplast, mitochondrial, and eukaryotic or that did not match any taxonomic phylum were excluded from the dataset. Samples that had less than 5,000 reads were removed at this stage. In order to merge studies from different sequence runs and primer sets, we grouped ASVs at the genus level before merging all studies into a single dataset in phyloseq v.1.36 (McMurdie and Holmes, 2013).

Microbial community assembly of the root microbiome was analyzed by performing a non-metric multidimensional scaling (NMDS) ordination utilizing the Bray-Curtis dissimilarity distance. Multivariate variation of the Bray-Curtis dissimilarity matrix was partitioned to ecosystem type, and compartment (soil/sediment, rhizosphere, and root), based on a PERMANOVA with 999 permutations performed in vegan v. 2.5.7 (Oksanen et al., 2013). Finally, putative function for sulfate reduction and sulfur oxidation was assigned based on homology at the genus level as in Rolando et al. (2022).

## Data availability

Raw metagenomic and metatranscriptomic sequences are deposited in the BioProject database (http://ncbi.nlm.nih.gov/bioproject) under accessions PRJNA703972 and PRJNA950121, respectively. Metagenome-assembled genomes are found in the BioProject PRJNA703972. Acompanying metadata and custom scripts used in the present study are publicly available in Supplementary Information and in a Zenodo repository: https://doi.org/10.5281/zenodo.7883423

## Acknowledgements

This work was supported in part by an institutional grant (NA18OAR4170084) to the Georgia Sea Grant College Program from the National Sea Grant Office, National Oceanic and Atmospheric Administration, US Department of Commerce, and by a grant from the National Science Foundation (DEB 1754756). Any opinions, findings, and conclusions or recommendations expressed in this material are those of the authors and do not necessarily reflect the views of the National Science Foundation.

## Author contributions

Conceptualization: J.L.R., M.K., J.T.M., and J.E.K. Field sampling and analysis: J.L.R., M.K., T.S., Y.L., and J.E.K. Bioinformatic analysis: J.L.R., M.K., P.P., R.C., and K.T.K. Funding acquisition: J.E.K. Writing—original draft: J.L.R. with inputs from all authors.

## Ethics declarations

The authors declare no competing interests.

---

**Supplementary Figure S1.**
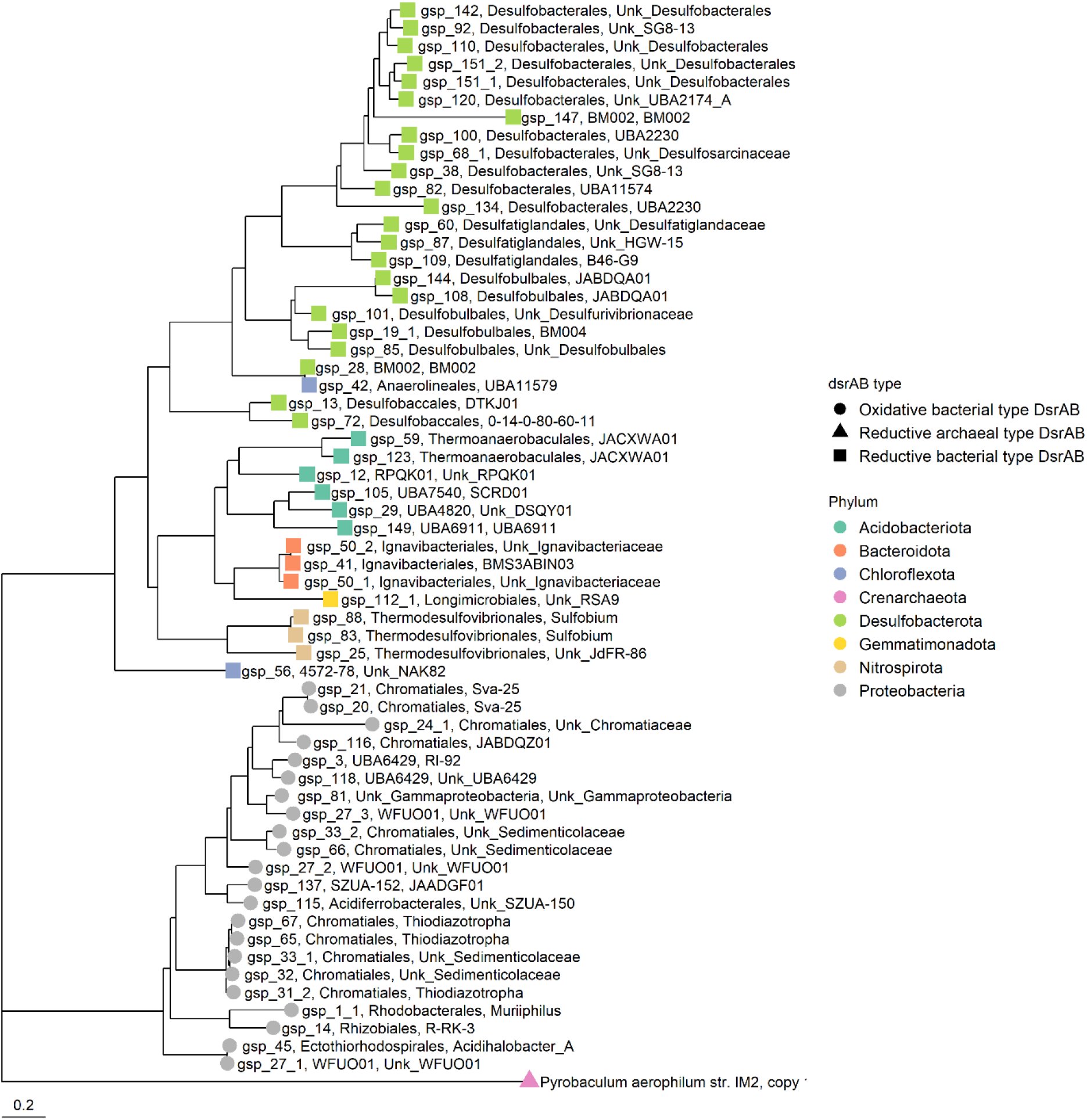
dsrAB phylogenetic tree of genes retrieved from metagenome-assembled genomes (MAGs). Genes were aligned to a reference database and tree was constructed with FastTree v.2.1.11. Oxidative and reductive types of the dsrAB gene were inferred based on phylogenetic placement in the constructed tree. Archean *Pyrobaculum aerophilum* was used as an outgroup.

**Supplementary Figure S2.**
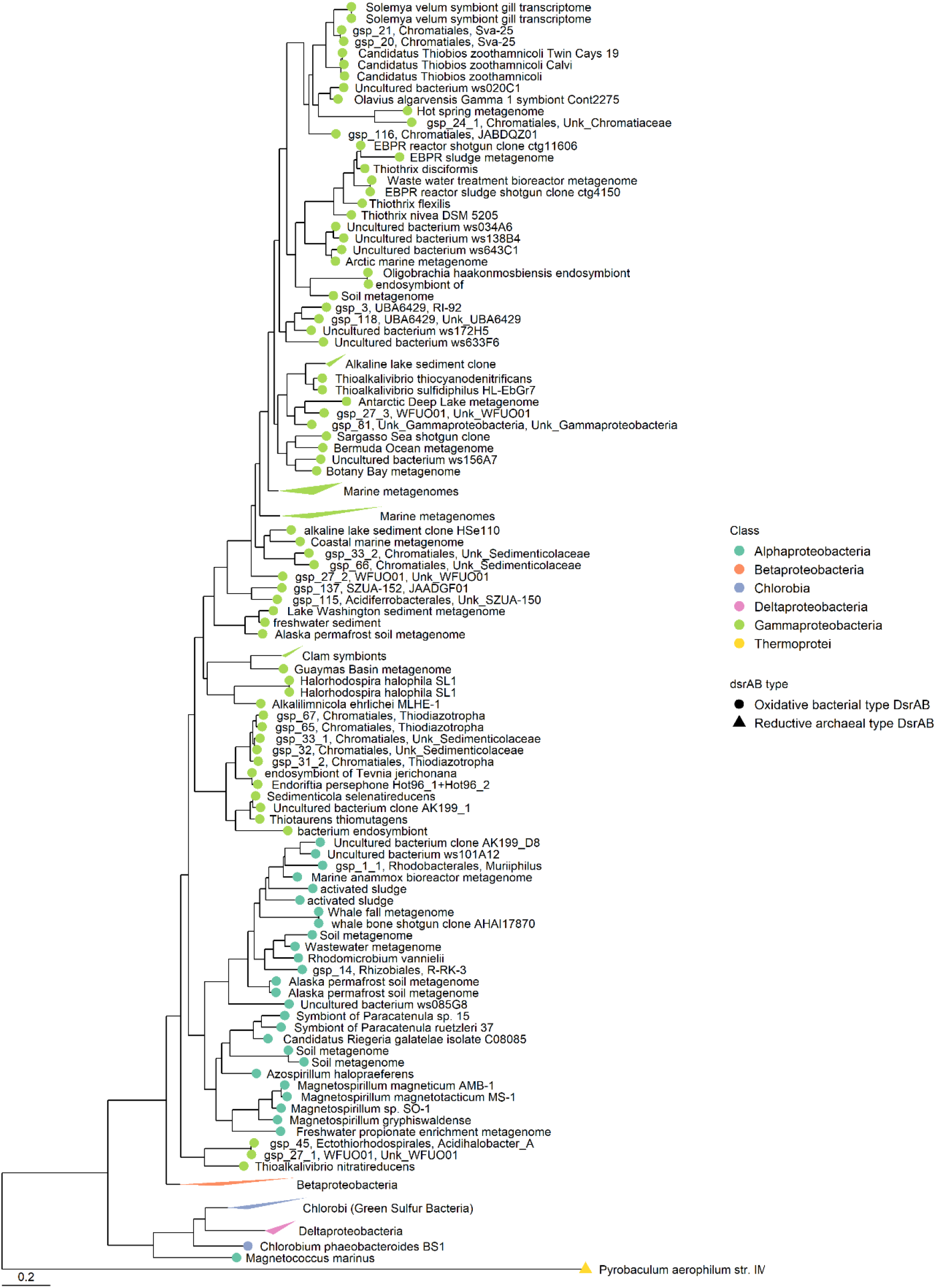
Phylogenetic tree of the oxidative dsrAB type from metagenome assembled genomes (MAGs) aligned to a reference alignment database (Müller et al., 2015). Archean *Pyrobaculum aerophilum* was used as an outgroup of the phylogenetic tree.

**Supplementary Figure S3.**
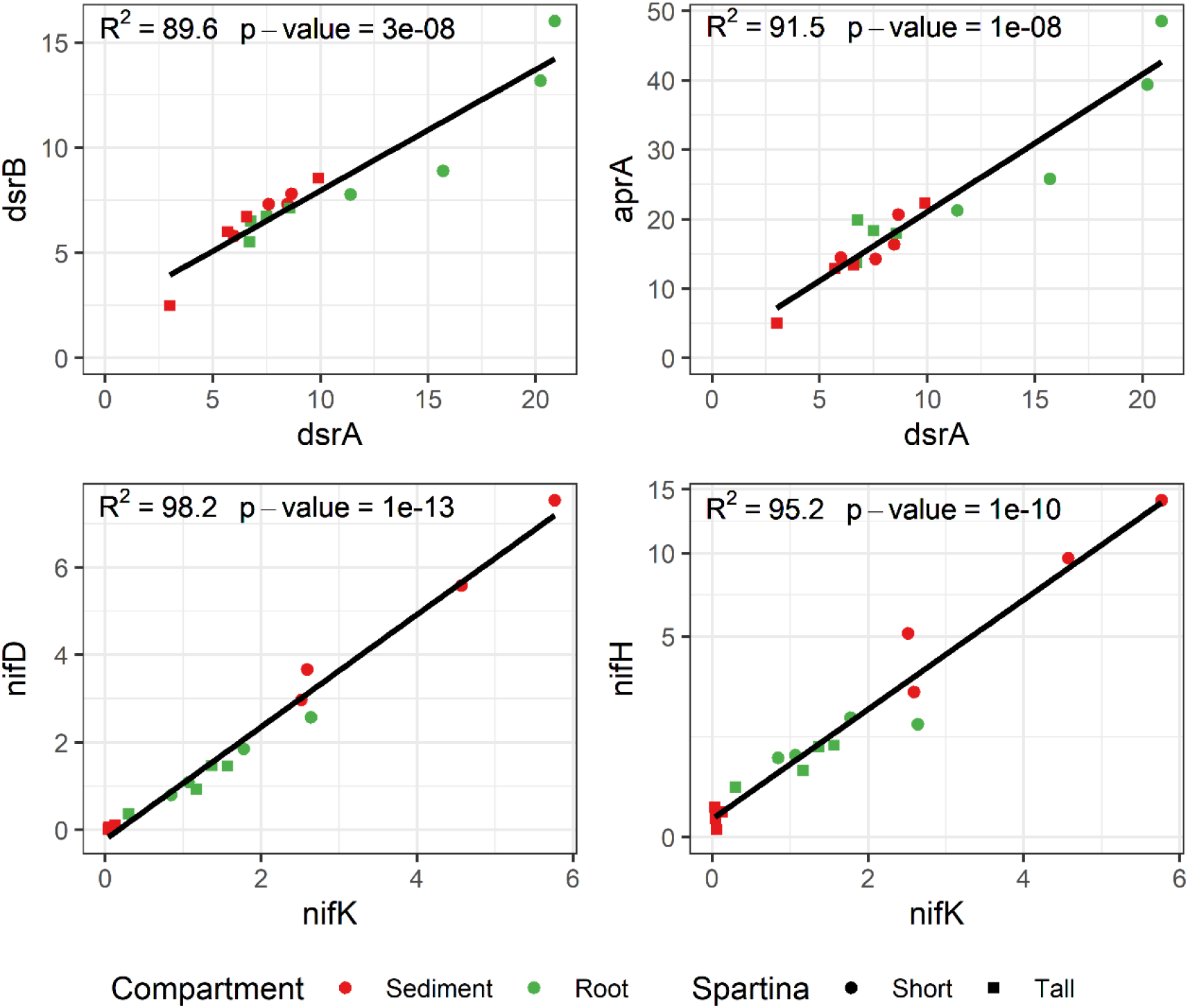
Gene expression of the nitrogenase (nifD, nifK, and nifH) and dissimilatory sulfite reductase genes (dsrA, dsrB, and aprA) are highly correlated among their own metabolic pathways.

**Supplementary Figure S4.**
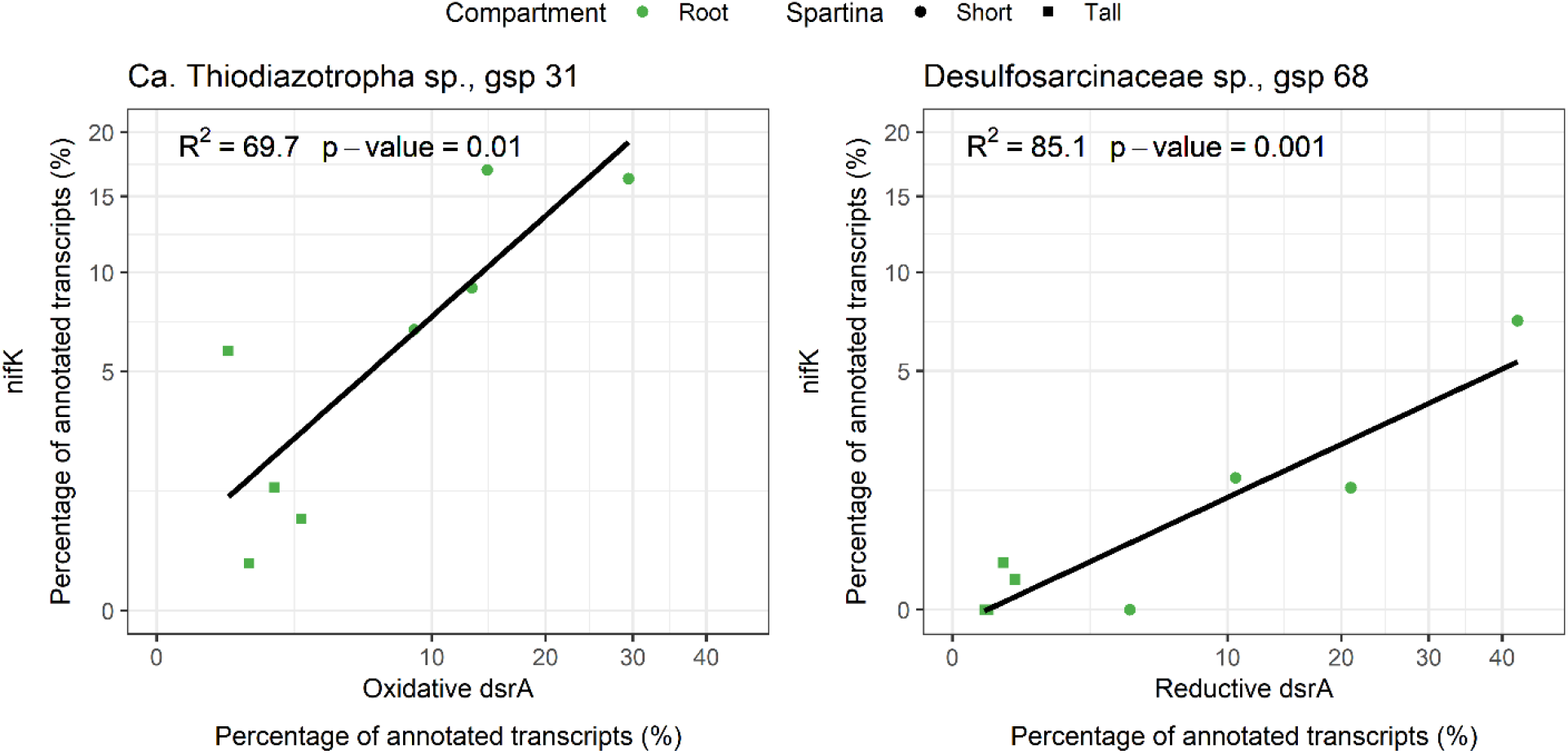
Relation between the gene expression of the *Ca.* Thiodiazotropha sp. (gsp 31) oxidative, and *Desulfosarcinaceae* sp. (gsp. 68) reductive dsrA gene and the nitrogenase gene nifK in *Spartina alterniflora* root samples. Gene expression measured as the percentage of transcripts mapping to each genomospecies from the totality of functionally annotated short read transcripts.

**Supplementary Figure S5.**
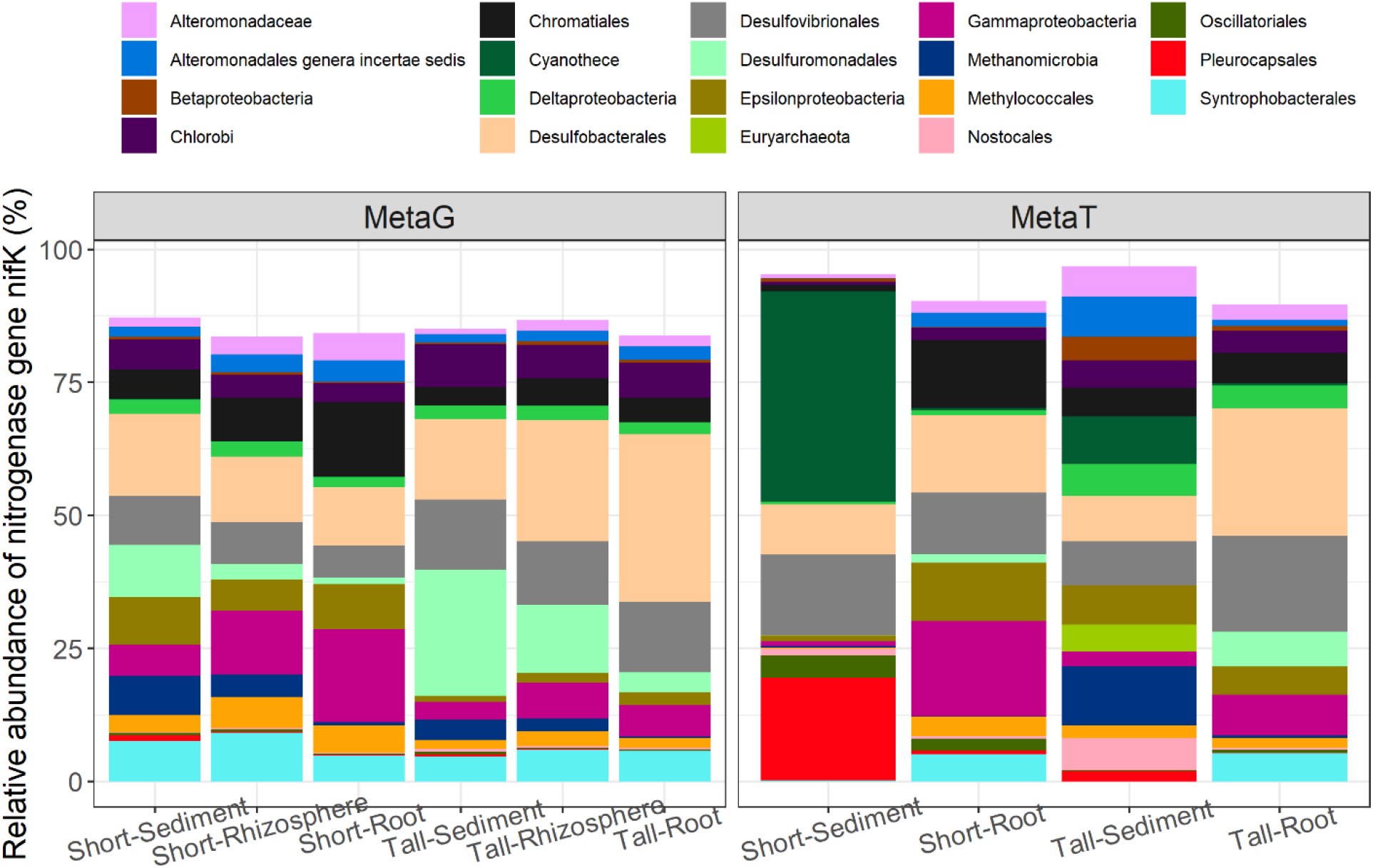
Gene and transcript relative abundance of the nitrogenase gene (nifK) partitioned at the finest taxonomic level as predicted by eggnog-mapper per microbiome compartment and *Spartina alterniflora* phenotype.

**Supplementary Figure S6.**
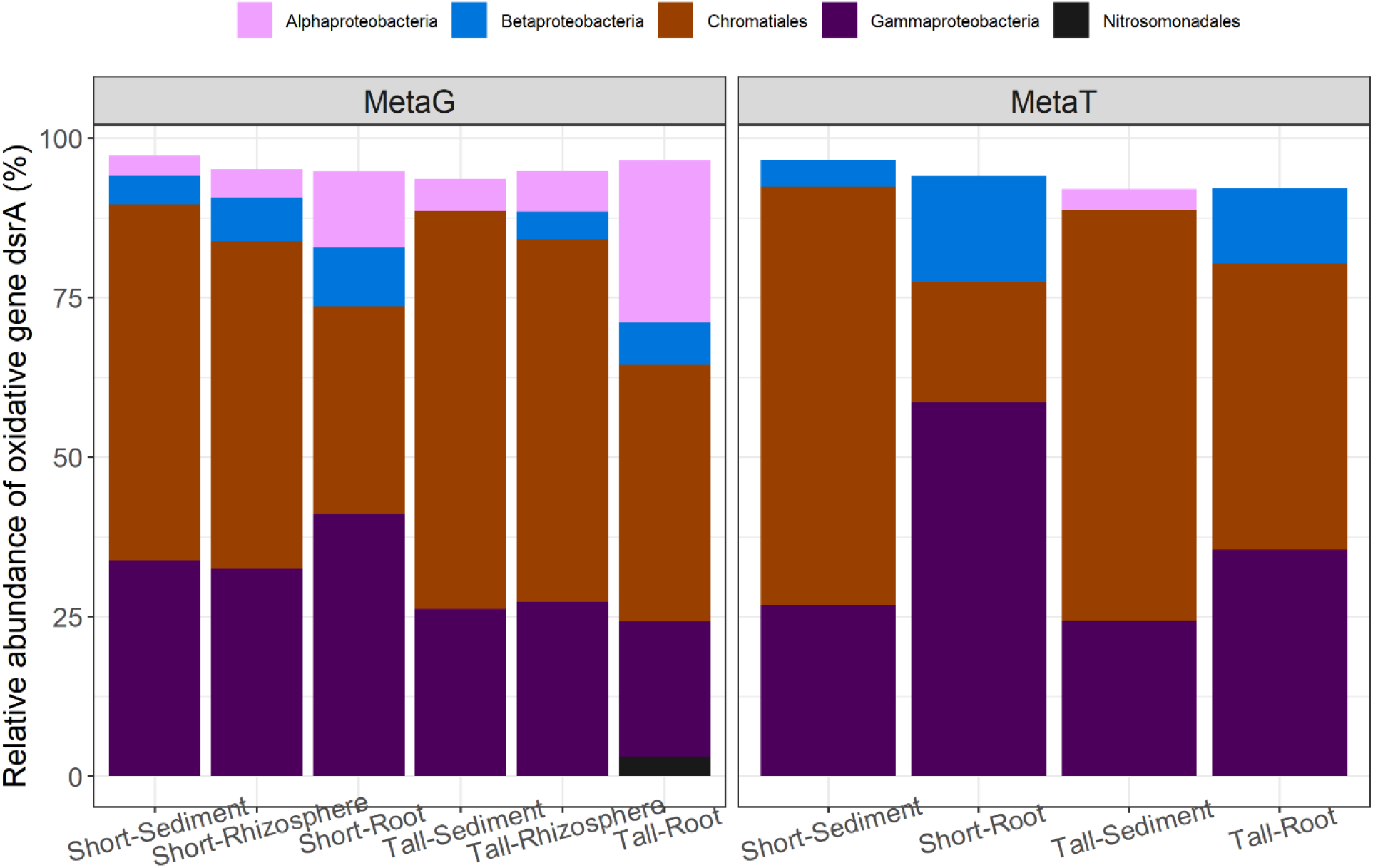
Gene and transcript relative abundance of the oxidative dsrA gene partitioned at the finest taxonomic level as predicted by eggnog-mapper per microbiome compartment and *Spartina alterniflora* phenotype.

**Supplementary Figure S7.**
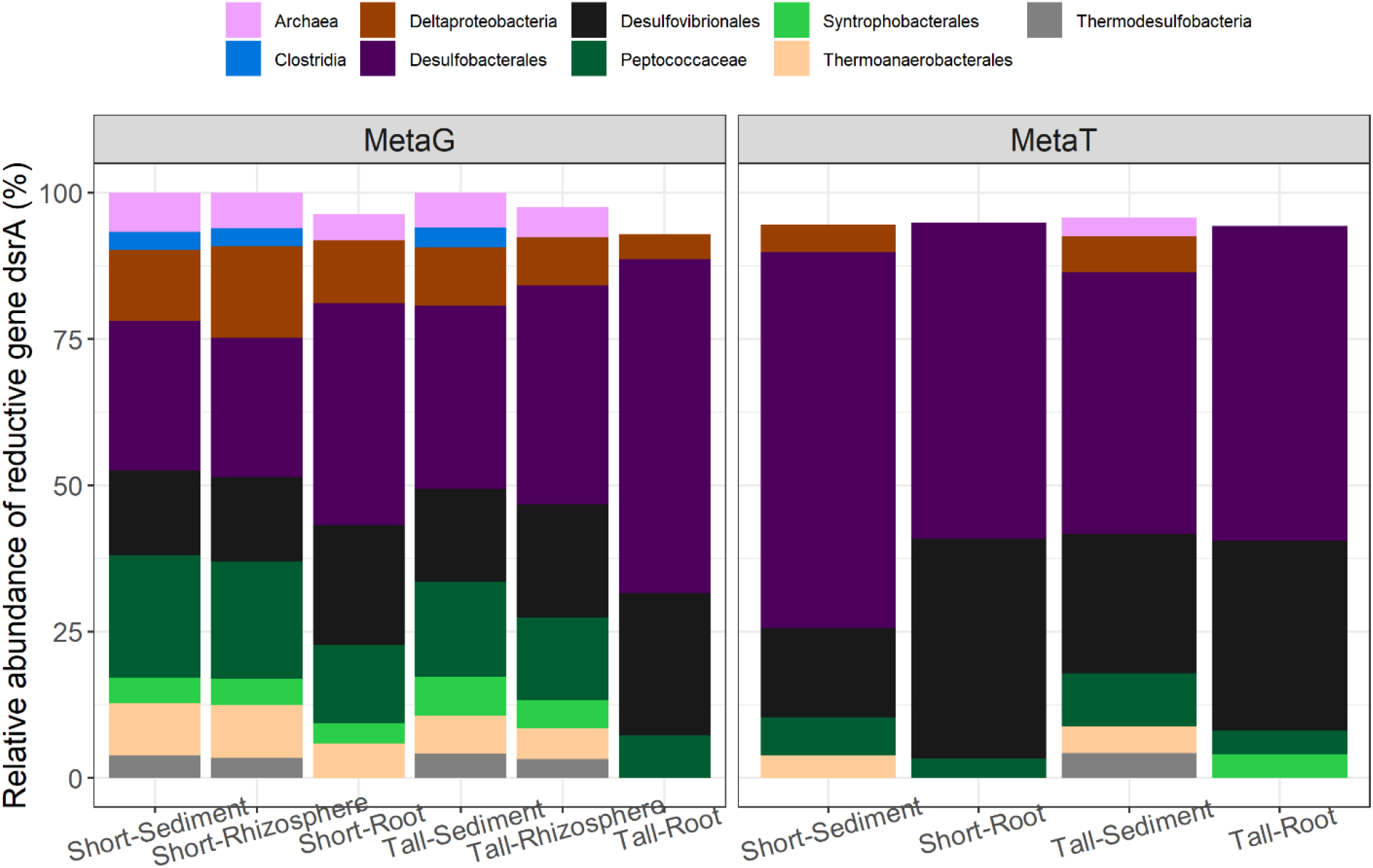
Gene and transcript relative abundance of the reductive dsrA gene partitioned at the finest taxonomic level as predicted by eggnog-mapper per microbiome compartment and *Spartina alterniflora* phenotype.

**Supplementary Figure S8.**
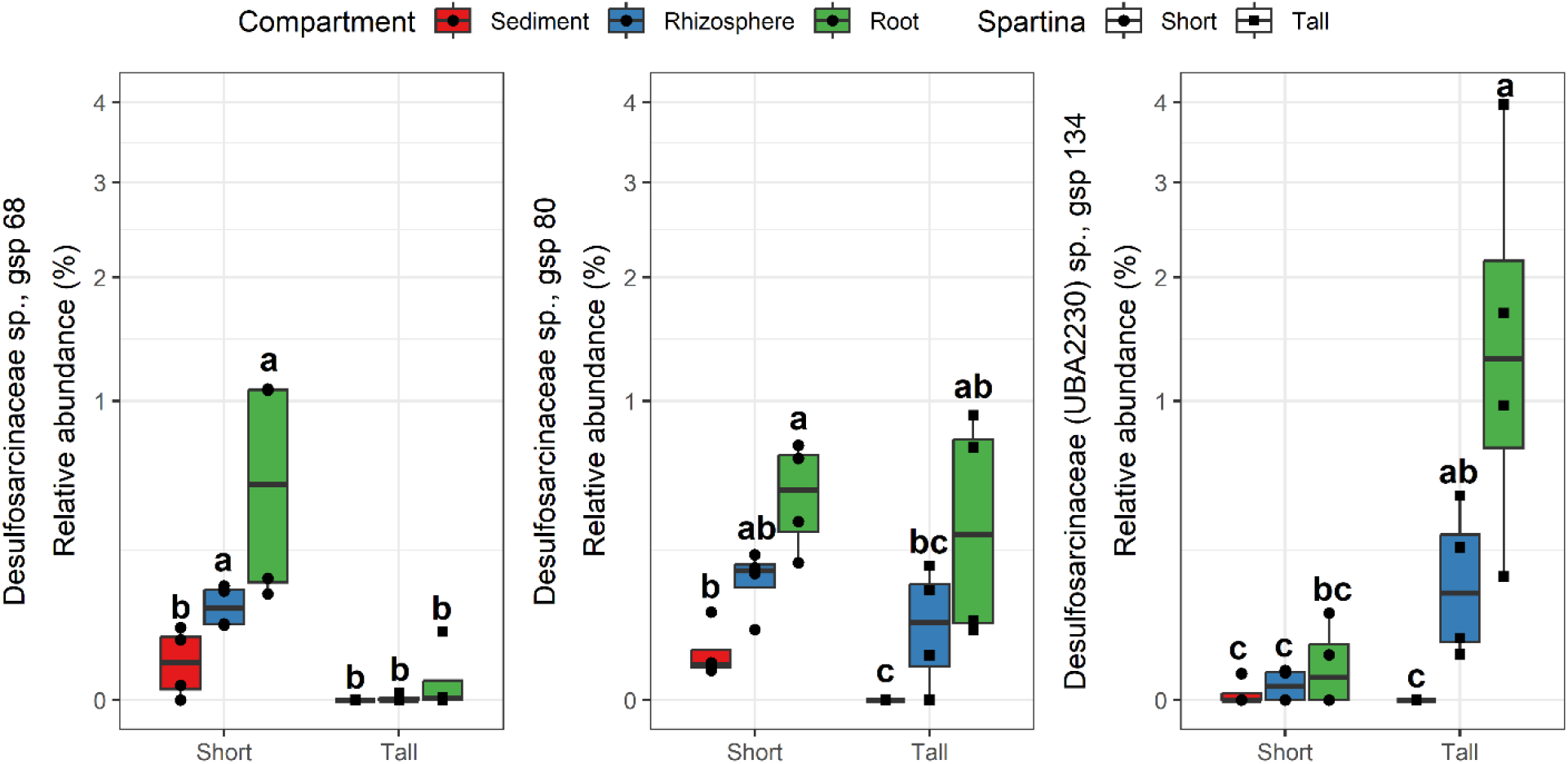
Relative abundance of selected metagenome assembled genomes (MAGs) from the *Desulfosarcinaceae* family across *Spartina alterniflora* phenotype and sampled compartment. Relative abundance was calculated at the DNA-level based on average coverage per position in metagenomic libraries. Different letter indicate statistical difference based on pairwise Mann-Whitney tests (p-value < 0.5).

**Supplementary Figure S9.**
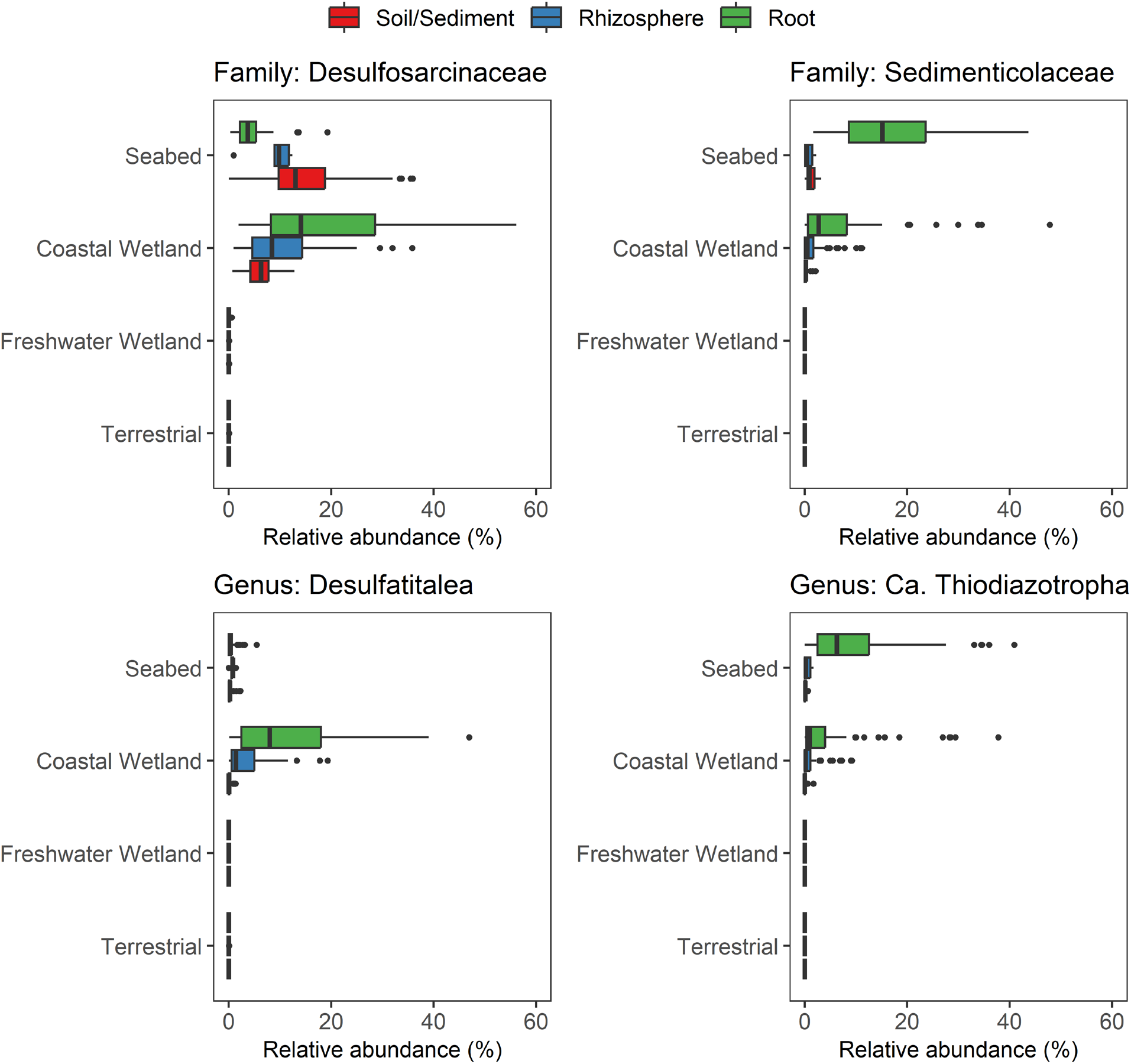
Relative abundance of amplicon sequence variants (ASVs) classified as members of the *Desulfosarcinaceae* and *Sedimenticolaceae* families, and the *Desulfatitalea* and *Ca.* Thiodiazotropha genera across soil/sediment, rhizosphere, and root samples from a seabed, coastal wetland, freshwater wetland and terrestrial ecosystems dataset.

**Supplementary Figure S10.**
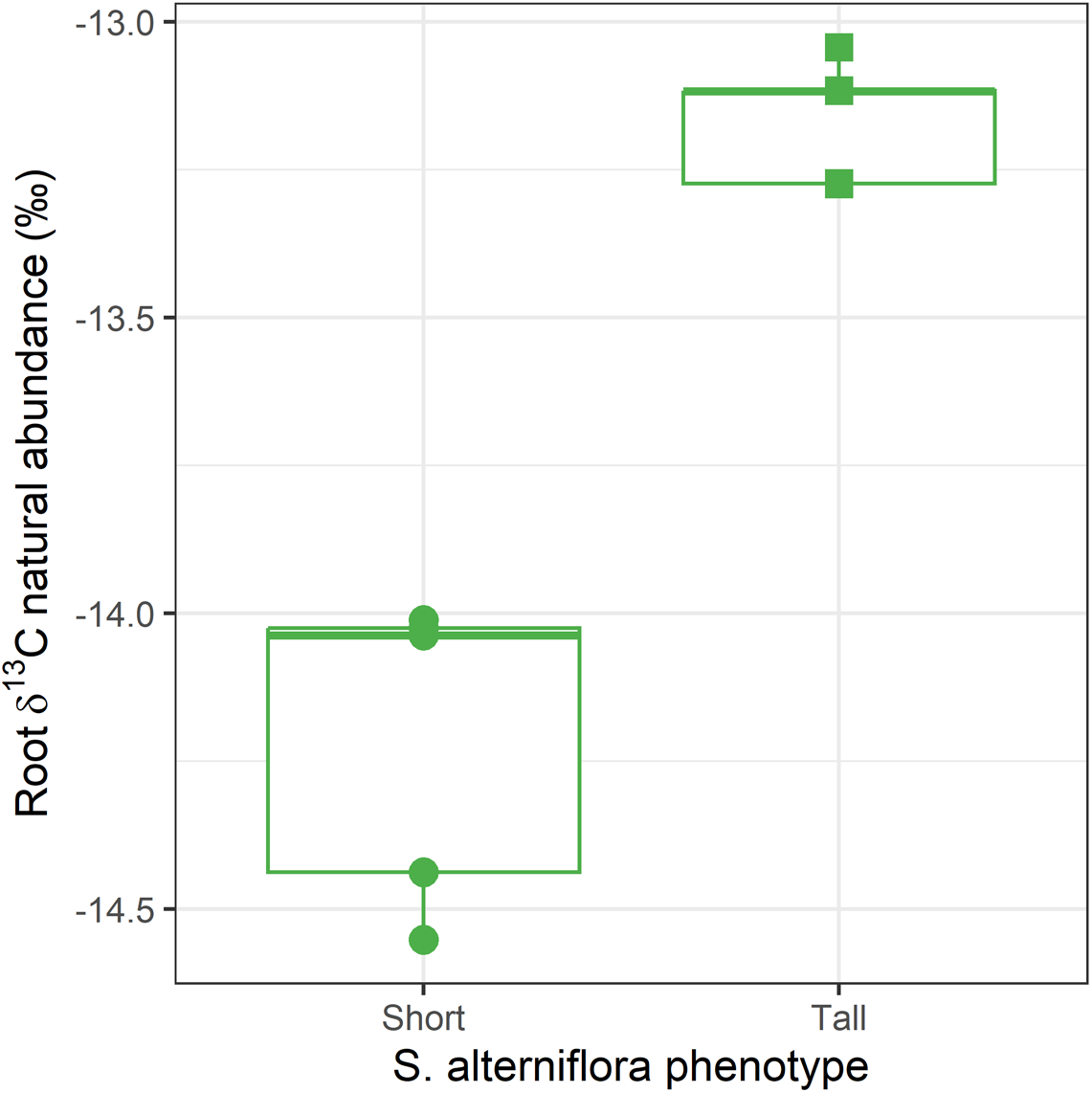
Root carbon ^13^C isotopic natural abundance by *Spartina alterniflora* phenotype. ^13^C natural abundance was expressed as the per mille (‰) deviation from the Pee Dee Belemnite standard (PDB) ^13^C:^12^C ratio (δ^13^C).

## Notes

### Competing Interest Statement

The authors have declared no competing interest.

### Summary of Updates

Fixed an error on author's order

